# A ribosome-associating chaperone mediates GTP-driven vectorial folding of nascent eEF1A

**DOI:** 10.1101/2024.02.22.581594

**Authors:** Ibrahim M. Sabbarini, Dvir Reif, Kibum Park, Alexander J. McQuown, Anjali R. Nelliat, Charlotte Trejtnar, Volker Dötsch, Eugene I. Shakhnovich, Andrew W. Murray, Vladimir Denic

## Abstract

Eukaryotic translation elongation factor 1A (eEF1A) is a highly abundant, multi-domain GTPase. Post-translational steps essential for eEF1A biogenesis are carried out by bespoke chaperones but co-translational mechanisms tailored to eEF1A folding remain unexplored. Here, we find that the N-terminal, GTP-binding domain of eEF1A is prone to co-translational misfolding and using computational approaches, yeast genetics, and microscopy analysis, we identify the conserved yet uncharacterized yeast protein Ypl225w as a chaperone dedicated to solving this problem. Proteomics and biochemical reconstitution reveal that Ypl225w’s interaction with ribosomal eEF1A nascent chains depends on additional binding of Ypl225w to the UBA domain of nascent polypeptide-associated complex (NAC). Lastly, we show by orthogonal chemical genetics that Ypl225w primes eEF1A nascent chains for their subsequent binding to GTP and release from Ypl225w. Our work establishes eEF1A as a model system for chaperone-dependent co-translational folding and unveils a novel mechanism for GTP-driven folding on the ribosome.

## Main

Protein folding across all three kingdoms of life is guided by a variety of chaperones, many of which recognize their clients as nascent chains emerging from the ribosome exit tunnel^1^. In eukaryotes, the major burden of co-translational folding falls on heat shock protein 70 (Hsp70) paralogs and their co-chaperones; additional contributions critical to general protein folding come from the Hsp90 system, as well as the TRiC/CCT chaperonin which works to fold tubulin, actin, and many other proteins with complex topologies^2^.

How does the eukaryotic ribosome choreograph co-translational chaperones and nascent chain targeting/processing factors as they all vie for the limited surface area surrounding the nascent chain exit site^3^? One body of work has shown that the ribosome-associated complex (RAC) facilitates Hsp70 activation and binding to specific nascent chain sites soon after they emerge from the exit tunnel^4–7^. The translating ribosome can also dynamically recruit the highly abundant nascent polypeptide-associated complex (NAC) to several distinct regions surrounding the exit site. Here, NAC functions as a gatekeeper for sorting nascent chains towards the signal recognition particle (SRP) or a peptidase for N-terminal methionine excision^8,9^. By contrast, the mechanism by which NAC assists co-translational protein folding remains poorly understood^10^.

Recently, we found that Zpr1 and its co-chaperone Aim29 mediate de novo folding of the highly abundant and essential eukaryotic translation elongation factor 1A (eEF1A)^11,12^. eEF1A, a GTPase, delivers amino acid-charged tRNAs to the translating ribosome and is recycled by a regulated G protein cycle^13^. A unique aspect of the Zpr1-Aim29 folding mechanism is its reliance on conformational changes driven by GTP hydrolysis of its client. This process occurs post-translationally and is likely preceded by individual folding of the three eEF1A domains: the N-terminal GTP binding domain (domain I, DI) and the two C-terminal β-barrel domains (DII and DIII). Structure-guided, loss-of-function mutations in Zpr1 or Aim29 lead to growth arrest or slow growth dependent on the degree to which eEF1A misfolding drives induction of the heat shock response (HSR) or activation of the integrated stress response (ISR) via reduction in eEF1A levels due to proteasomal degradation.

We considered the possibility of additional eEF1A folding factors restricted to the co-translational stages of eEF1A biogenesis. This notion came partially from a single-molecule analysis of co-translational folding of EF-G, a prokaryotic GTPase that mediates ribosome translocation during elongation^14,15^. In this study, co-translational misfolding between the nascent N-terminal GTP-binding domain (DI) and the ensuing β-barrel domain II (DII) was suppressed by Trigger Factor, a ribosome-associating, ATP-independent chaperone unique to prokaryotes. Owing to the sequence and structural similarity between EF-G DI/DII and the corresponding eEF1A domains, we wondered if protein folding factors specific to eukaryotes play an analogous role during eEF1A biogenesis.

Indeed, in this study, we discover that the conserved protein Ypl225w acts as a chaperone tailored to the co-translational folding of eEF1A DI. We show that ribosomal recruitment of Ypl225w by NAC enables Ypl225w to associate with eEF1A nascent chains and thereby prime DI for GTP binding. The subsequent GTP binding step enables eEF1A DI folding concomitant with Ypl225w recycling. Our work reveals the first example of an ATP-independent chaperone system on the ribosome that co-opts the nucleotide binding of a nascent G protein client into a folding switch.

## Results

### Computational modeling identifies Ypl225w as an eEF1A chaperone candidate

To search for new eEF1A chaperones, we used a version of AlphaPulldown^16^ (a derivative of AlphaFold) specific for the budding yeast *Saccharomyces cerevisiae*. In this screening approach, eEF1A’s sequence served as the bait against a library of “prey” sequences consisting of yeast proteins conserved across eukaryotes. We filtered interactions based on structures with an ipTM (interface pTM, a measure of structure confidence) greater than 0.8 (see Methods) (Fig. 1a). The top hits were two bona fide eEF1A physical and functional interactors (Efb1 and Efm4)^17,18^, as well as the uncharacterized but conserved protein Ypl225w. The structural model of the Ypl225w•eEF1A interaction predicts a high confidence interface (ipTM = 0.82) in which the N-terminal α-helix (NaH) of Ypl225w inserts itself into the GTP-binding, switch regions of eEF1A DI, in a manner that is largely unaffected by the additional presence of DII and DIII (Fig. 1b and Extended Data Fig. 1a-b).

**Fig. 1.**
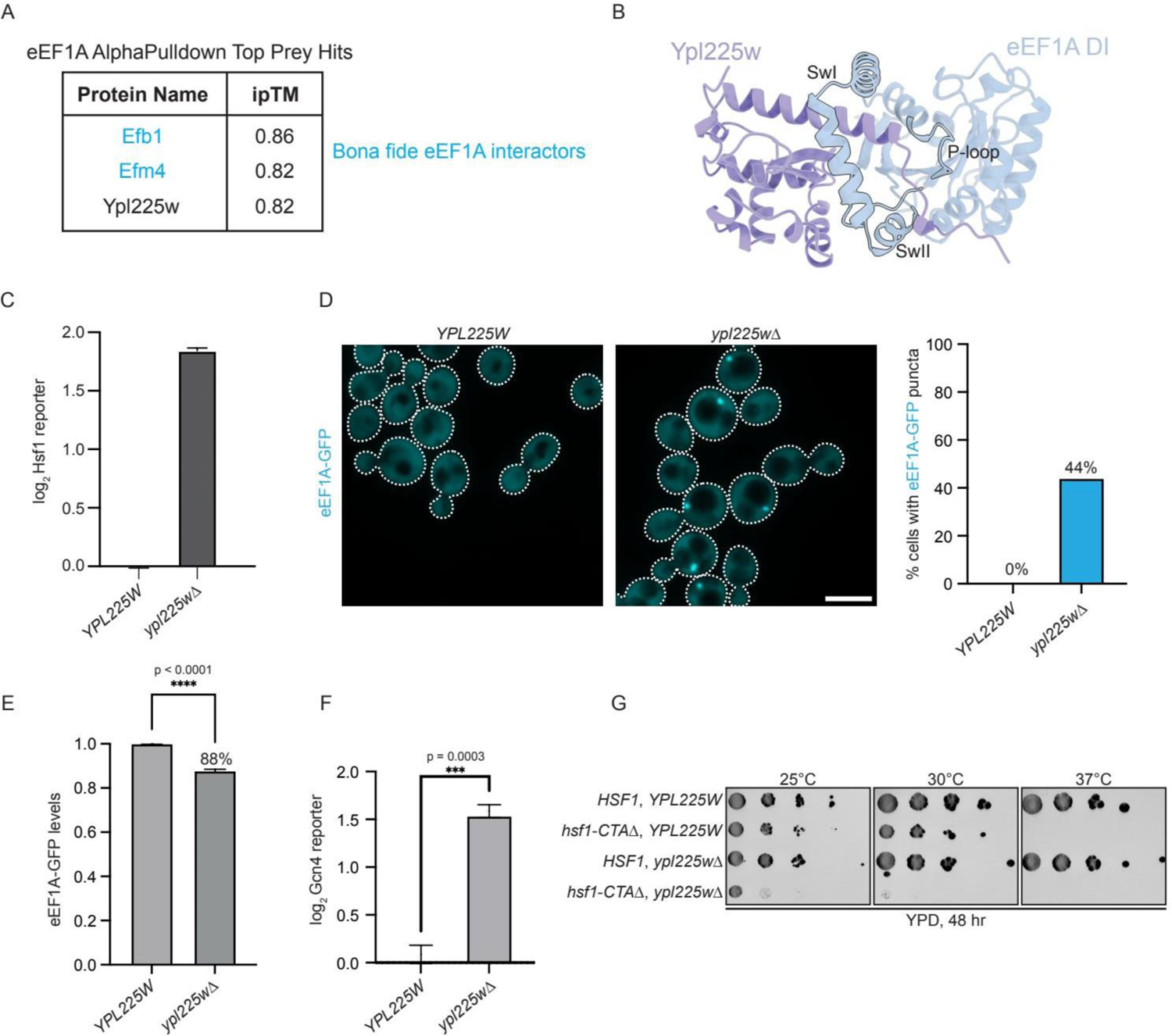
Loss of the eEF1A chaperone candidate Ypl225w elicits the hallmarks of eEF1A biogenesis defects *in vivo*. (A) Table showing the top three hits (ipTM > 0.8) from an AlphaPulldown screen that utilized eEF1A’s sequence as bait. Bona fide eEF1A interactors are highlighted in blue. (B) ColabFold model showing eEF1A DI (light blue) bound to Ypl225w (light purple). The Switch I (SwI), Switch II (SwII), and P-loop regions of eEF1A DI are outlined in black. (C) The indicated strains carrying the Hsf1 reporter (*4xHSE-YFP*) were analyzed by flow cytometry. Bar graphs show median YFP intensity values normalized to both cell size (side scatter area) and the average median YFP intensity of wild-type (*YPL225W*) cells. Error bars represent the standard deviation from three biological replicates. (D) Wild-type (*YPL225W*) and *ypl225w*Δ strains expressing endogenously-tagged eEF1A-GFP were imaged by confocal microscopy. Shown are representative micrographs normalized to the same intensity, with cells outlined. Scale bar represents 2.5 µm. 100 cells were quantified for each sample (right panel). (E) Wild-type (*YPL225W*) and *ypl225w*Δ strains expressing endogenously-tagged eEF1A-GFP were analyzed by flow cytometry as described in panel C. Shown are bar graphs of median GFP intensity values normalized to both cell size and the average median GFP intensity of wild-type cells. Measurements were compared by an unpaired, two-tailed t-test. **** p < 0.0001. (F) The indicated strains carrying a translational reporter of Gcn4 synthesis (upstream ORFs from *GCN4* driving GFP expression) were analyzed by flow cytometry. Bar graphs and measurements of reporter activity were generated and analyzed as in panel C. Measurements were compared by an unpaired, two-tailed t-test. *** p < 0.001. (G) Strains bearing the indicated genotypes were grown to mid-log phase (OD_600_ = 0.4) in YPD media, serially diluted by a factor of 10, and spotted onto YPD plates prior to being incubated at the indicated temperatures. Plates were imaged after 48 hours.

We looked for additional computational evidence that Ypl225w and eEF1A constitute a chaperone-client relationship using co-translational folding simulations^19–21^. This approach revealed that the region of eEF1A predicted to interact with Ypl225w (amino acids 1-72) is prone to forming a kinetically-trapped misfolded intermediate (Extended Data Fig. 1c). To define the sequence specificity of these findings, we analyzed a highly similar region in Sup35, a translation termination GTPase in *S. cerevisiae* and a closely-related paralog of eEF1A^22^. Strikingly, the predicted co-translational folding pathway of this Sup35 region was populated only by native-like folding intermediates (Extended Data Fig. 1c). Additionally, ColabFold modeling found no strong evidence of an interaction between Ypl225w and Sup35, supporting the hypothesis that Ypl225w and eEF1A are a selective chaperone-client pair (Extended Data Fig. 1d).

Two findings in the literature are consistent with Ypl225w’s potential role in co-translational eEF1A folding: 1) proteomic analysis of purified yeast ribosomes identified Ypl225w as an associated factor^23^, and 2) a genome-wide screen for mutations that chronically induce an HSR reporter ranked *ypl225w*Δ as the 12^th^ highest hit^24^. When the latter study was completed, deletion of *AIM29*, now known to encode a chaperone for eEF1A, was the only other uncharacterized gene that ranked higher than *ypl225w*Δ.

In summary, our computational modeling and data mining led us to formulate a structure-based hypothesis in which Ypl225w uses its NaH to chaperone early eEF1A nascent chains emerging from the ribosome.

### *ypl225w*Δ cells display all the hallmarks of defective eEF1A biogenesis

We began by examining *ypl225w*Δ cells for signs of disrupted eEF1A biogenesis that we saw previously in cells lacking Zpr1 or Aim29^11,12^. First, we confirmed that loss of Ypl225w elevated basal Hsf1 activity by utilizing the same HSR reporter used in the aforementioned genome-wide study^24–26^ (Fig. 1c). Second, we looked for evidence of eEF1A-GFP aggregates in *ypl225w*Δ cells and observed fluorescent punctae in nearly half of the imaged cells (44%; Fig. 1d). Third, we detected a small but significant reduction in eEF1A protein levels (12% relative to WT cells) by flow cytometry analysis of eEF1A-GFP fluorescence (Fig. 1e). These latter phenotypes – aggregation and reduced abundance – are the expected outcomes of eEF1A misfolding leading to protein aggregation or degradation^12^. Lastly, *ypl225w*Δ induced a translational reporter of Gcn4 synthesis, a sign of the integrated stress response, which is expected from the reduction in eEF1A level (Fig. 1f)^12^.

*ypl225w*Δ cells grow normally under standard conditions and the eEF1A biogenesis defects reported above are milder than those caused by mutations disrupting Zpr1 or Aim29 function (Extended Data Fig. 1e). However, when we replaced wild-type Hsf1 with a C-terminally truncated allele that cannot induce a heat shock response at elevated temperatures^27^, we observed a strong synthetic growth defect in *ypl225w*Δ cells (Fig. 1g). These data suggest that an adaptive HSR in *ypl225w*Δ cells partially restores eEF1A proteostasis towards its original set-point. Finally, we tested whether the ability of Ypl225w to maintain proteostasis was conserved across two diverged yeast species (>500 MYA) and found that heterologous expression of the *Schizosaccharomyces pombe* Ypl225w ortholog (*SPBC3E7.07c*) fully suppressed the HSR in *ypl225w*Δ cells (Extended Data Fig. 1f-g). Collectively, these findings reveal that Ypl225w has a conserved but poorly defined function in eEF1A biogenesis.

### Ypl225w is a *bona fide* eEF1A chaperone

To test Ypl225w’s biochemical activity as a co-translational eEF1A chaperone, we turned to our *in vitro* translation system for studying eEF1A biogenesis and found several lines of supporting evidence for Ypl225w’s role as a chaperone^11,12^. First, eEF1A synthesized in *ypl225w*Δ extracts acquired only ∼20-30% of its trypsin resistance relative to the wild-type control (Fig. 2a). Second, co-translationally adding back pure recombinant Ypl225w-3xFLAG restored eEF1A folding back to wild-type levels (Fig. 2b). Third, the reduced ability of Ypl225w-3xFLAG, added after protein synthesis, to restore eEF1A folding demonstrated that Ypl225w was more effective co-translationally (Fig. 2c). Finally, in agreement with our computational modeling, we observed that the ability of newly-synthesized Sup35 to acquire trypsin resistance is independent of Ypl225w (Extended Data Fig. 2a).

**Fig. 2.**
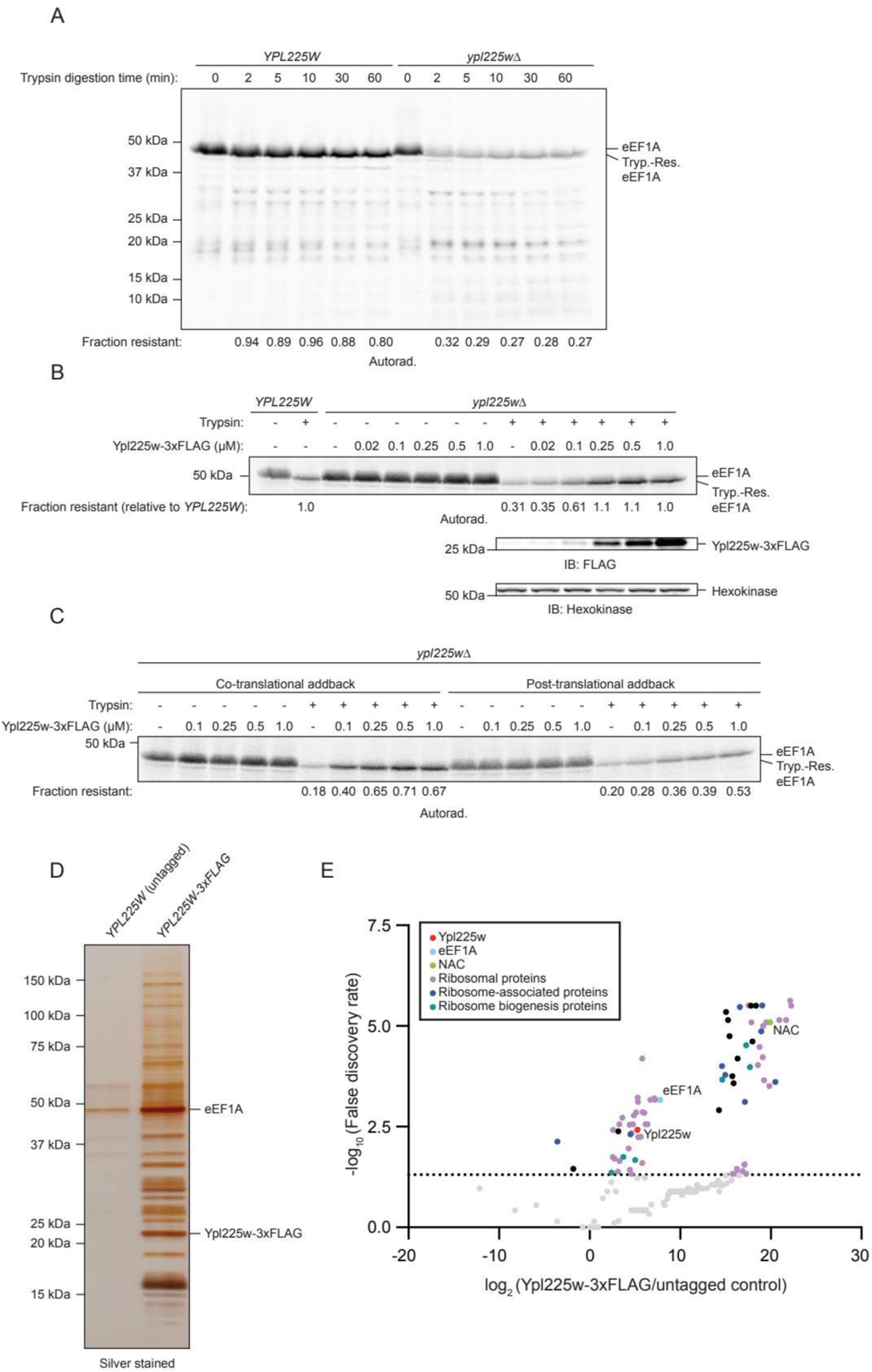
Ypl225w is a bona fide eEF1A co-translational chaperone. (A) Trypsin resistance of ^35^S-methionine-radiolabeled eEF1A was determined following *in vitro* translation (IVT) for 45 minutes in wild-type (*YPL225w*) or *ypl225w*Δ extracts and addition of 100 µg/mL cycloheximide (CHX) to terminate further protein synthesis. The ‘0’ time point represents the undigested sample. Following digestion for the indicated times, samples were resolved by SDS-PAGE and visualized by autoradiography. ‘Fraction resistant’ is the signal of near full-length, trypsin-resistant (Tryp.-Res.) eEF1A fragment divided by the total synthesized eEF1A in the undigested ‘0’ timepoint. (B) As in panel A but with all reactions analyzed after 5 minutes of trypsin digestion and with addition of the indicated amounts of recombinant Ypl225w-3xFLAG protein prior to IVT. Samples were also analyzed by immunoblotting (IB) with the indicated antibodies. ‘Fraction resistant’ is the signal of near full-length, trypsin-resistant (Tryp.-Res.) eEF1A fragment divided by the total synthesized eEF1A in the corresponding undigested lane normalized to the WT *YPL225W* signal. (C) As in panel B but with a translation time of 20 minutes. ‘Co-translational addback’ reactions were supplemented with Ypl225w-3xFLAG protein prior to translation initiation and immediately treated with trypsin post-CHX addition. ‘Post-translational addback’ reactions were only supplemented with Ypl225w-3xFLAG protein after CHX treatment and then incubated at room temperature for 20 minutes before trypsin treatment. (D) Affinity-purified samples from one replicate of the untagged (*YPL225W*) and *YPL225W-3xFLAG* samples that were used for mass spectrometry analysis shown here after SDS-PAGE followed by silver staining. (E) Mass spectrometry analysis of samples similar to the ones in panel D. The dotted line represents a false discovery rate at the p-value of 0.05. The inset within the volcano plot represents various categories of the enriched Ypl225w-3xFLAG interactors. The black dots represent interactors outside of the indicated categories.

### Ypl225w association with ribosomes is dependent on ongoing eEF1A synthesis

We hypothesized that Ypl225w’s previously noted ribosome association^23^ might reflect its preference for co-translational eEF1A clients. To first corroborate the ribosomal association, we immunoprecipitated Ypl225w-3xFLAG from cell extracts and quantitatively analyzed its interactors by SDS-PAGE and mass spectrometry (Fig. 2d-e and Supplementary Table 1: IP-MS data). Besides eEF1A, we detected numerous large and small ribosomal subunit proteins among the top hits (Fig. 2e and Supplementary Table 1: IP-MS data). In a complementary approach, we subjected extracts to sucrose gradient fractionation. Western blotting revealed a population of Ypl225w-3xFLAG in the heavy fractions containing polysomes, which we could eliminate by pre-treating extracts with EDTA, a condition that dissociates ribosomal subunits (Extended Data Fig. 2b).

Next, we asked if Ypl225w association with ribosomes depends on ongoing eEF1A biogenesis. For this purpose, we introduced Ypl225w-3xFLAG into cells engineered for eEF1A expression under the *GAL1* promoter (Tef-Off). As we have shown previously, Tef-Off cells grow normally in the presence of galactose but become terminally arrested following a prolonged period of promoter-shutoff in glucose-containing media^12^. In the current application of this system, we induced promoter shut-off for a shorter period of time that did not slow down cell growth but was in principle sufficient to clear pre-existing eEF1A mRNAs (average half-life ∼30 minutes)^28^. Consistent with the idea that cells at that point in the shut-off still have enough mature eEF1A to maintain translation, we observed a similar polysome to monosome ratio with that of control cells grown in galactose (Extended Data Fig. 2c). Strikingly, our brief Tef-Off promoter-shutoff period induced the disappearance of Ypl225w-3xFLAG from the heavy, polysome-containing fractions (Extended Data Fig. 2c). In sum, these findings confirm that Ypl225w associates with ribosomes and provide new evidence that this association depends on eEF1A nascent chains.

### Ypl225w is a co-translational chaperone for the GTP-binding domain of eEF1A

Our ColabFold model of the Ypl225w•eEF1A complex suggested that Ypl225w specifically chaperones the DI portion of eEF1A nascent chains. We found two additional lines of evidence supporting this hypothesis. First, we fused eEF1A DI to GFP and observed signs of protein aggregation and degradation in *ypl225w*Δ cells, phenotypes that were absent from the wild type (Fig. 3a-b and Extended Data Fig. 3a). By comparison, cells acutely depleted of Zpr1 do not form eEF1A DI-GFP aggregates^12^, further supporting Ypl225w’s DI-tailored role. This effect was selective to eEF1A as Sup35 DI-GFP revealed no comparable evidence of misfolding in *ypl225w*Δ cells (Extended Data Fig. 3a).

**Fig. 3.**
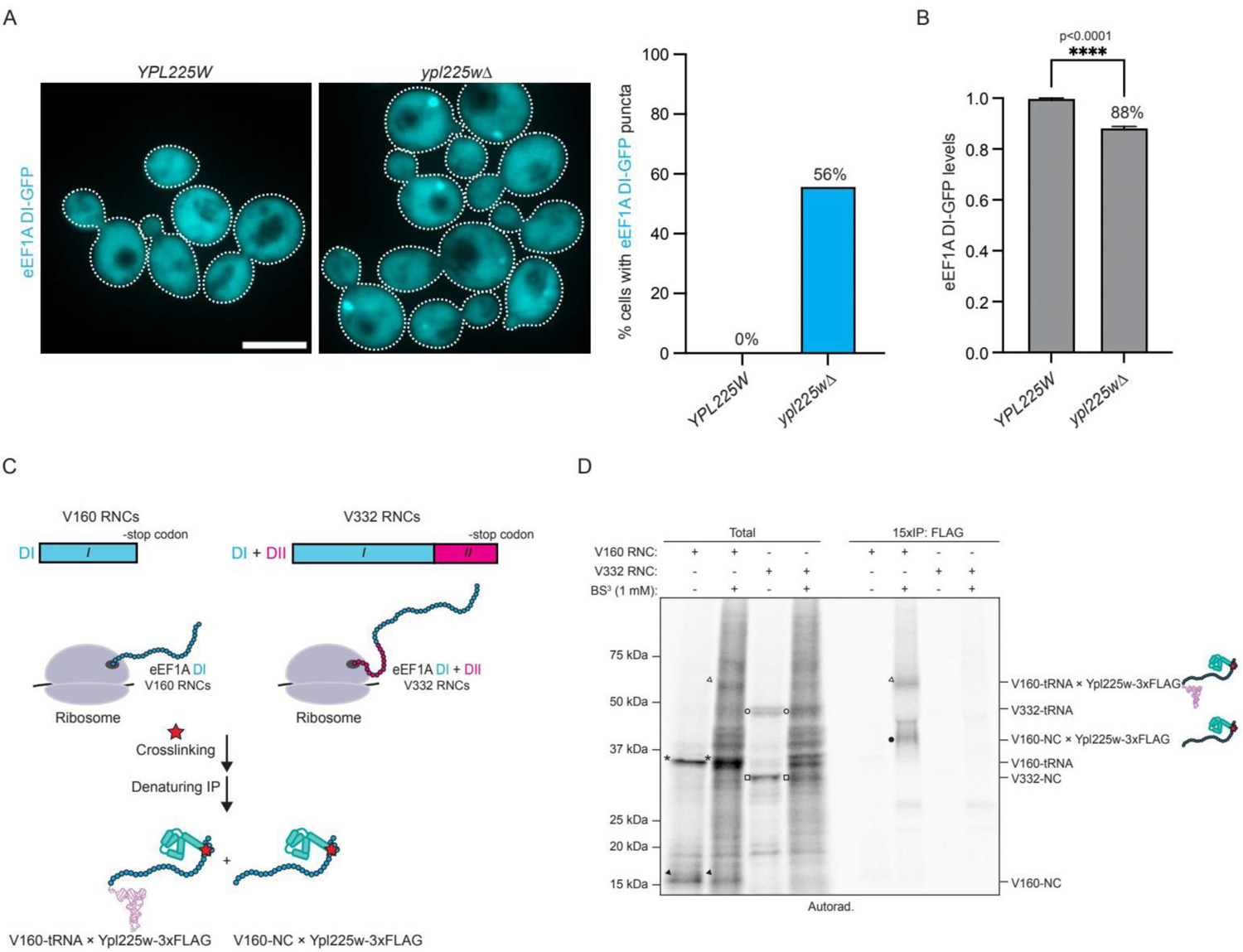
Ypl225w’s chaperone function is restricted to eEF1A DI. (A) WT (*YPL225W*) and *ypl225w*Δ strains expressing eEF1A DI-GFP driven by the *GAL1* promoter (integrated at the *URA3* locus) were imaged by confocal microscopy and analyzed as in Fig. 1d. Scale bar represents 2.5 µm. 100 cells were quantified for each sample (right panel). See STAR Methods for details of how expression of *GAL1* was induced. (B) Strains expressing eEF1A DI-GFP from panel A were analyzed via flow cytometry as described in Fig. 1e. Measurements were compared by an unpaired, two-tailed t-test. **** p < 0.0001. (C) Cartoon schematic of ribosome-nascent chain complexes (RNCs) and sample processing for RNC-cross-linking experiment in panel D. V160/V332 represent stalled RNCs of 160 or 332 amino acids in length stalled at a valine (V) codon. (D) IVT reactions from *ypl225w*Δ extracts supplemented with 1 µM Ypl225w-3xFLAG and nonstop eEF1A mRNAs (V160 or V332 RNCs) were translated for 15 minutes prior to cross-linking with 1 mM BS^3^. Samples were then subjected to denaturing FLAG immunoprecipitations (IPs) and analyzed by SDS-PAGE followed by autoradiography. Filled arrow: V160-NC, asterisk: V160-tRNA, open arrow: V160-tRNA x Ypl225w-3xFLAG, filled circle: V160-NC x Ypl225w-3xFLAG, open square: V332-NC, open circle: V332-tRNA.

In the second approach, we examined the “timing” of Ypl225w’s association with ribosome-nascent chain complexes (RNCs) using cell-free translation^29^. Here, we stabilized RNCs at the 3’ ends of two mRNAs encoding truncated versions of eEF1A but lacking the terminal stop codon (Fig. 3c). The shorter truncation, V160-RNC, represents a translation elongation intermediate in which only a portion of DI has been synthesized, while the longer truncation, V332-RNC proceeds to additionally synthesize DII. To specifically monitor Ypl225w-3xFLAG interaction with these RNCs, we used chemical cross-linking followed by denaturation and immunoprecipitation (Fig. 3c). This approach allowed us to detect cross-linked adducts between Ypl225w and V160-nascent chain (NC) but not V332-NC (Fig. 3d). We used RNase digestion to confirm that the higher molecular weight adduct corresponded to tRNA-conjugated V160-NC while the other species are deacylated products thereof, likely formed during sample processing (Extended Data Fig. 3b). These *in vitro* data strongly support a model in which Ypl225w interacts with nascent chains of eEF1A’s DI but completes its chaperone function as elongation proceeds further.

### NAC facilitates eEF1A folding *in vitro* by ribosomal recruitment of Ypl225w

In considering how Ypl225w might gain access to eEF1A RNCs, we noted the presence of the NAC on our list of interactors (Fig. 2e). The major form of yeast NAC comprises the Egd2 α subunit in complex with the Egd1 β subunit^30^. Our ColabFold modeling of Ypl225w’s interaction with NAC revealed a high confidence interface with the C-terminal ubiquitin-associated (UBA) domain of Egd2, which could accommodate the nearby Ypl225w-eEF1A DI interaction (Fig. 4a). Intriguingly, a similar interface between NAC UBA and the signal recognition particle (SRP) was recently resolved by CryoEM analysis of RNCs with an exposed ER signal sequence (Extended Data Fig. 4a)^9^. This interaction is critical for tethering SRP to the ribosome prior to receiving the ER signal sequence from NAC. Since NAC, to our knowledge, has not been previously shown to recruit folding factors to the ribosome, we decided to explore this possibility in the context of Ypl225w-mediated eEF1A folding.

**Fig. 4.**
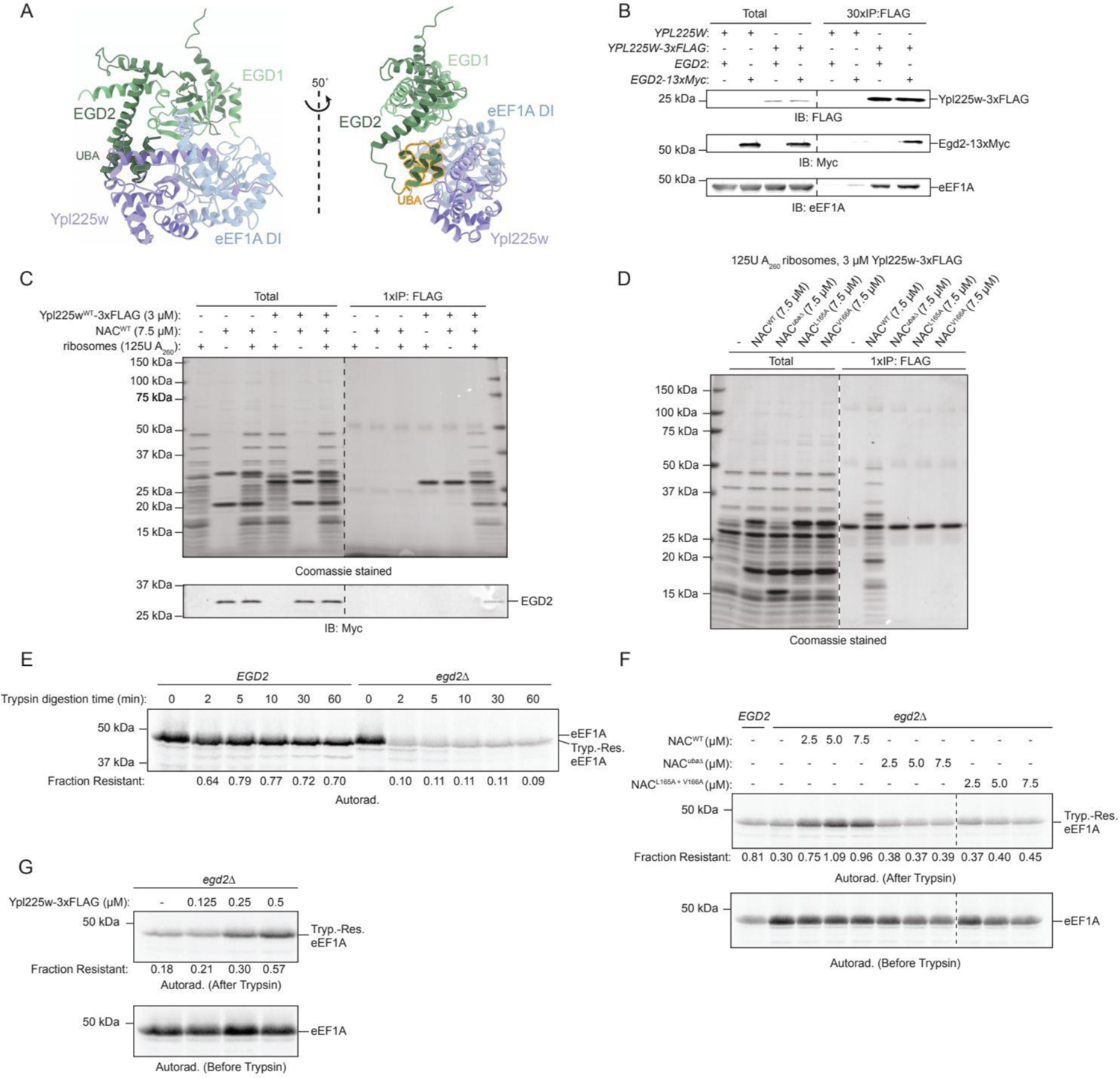
NAC facilitates eEF1A co-translational folding by recruiting Ypl225w to ribosomes. (A) ColabFold models of eEF1A DI (light blue) bound to Ypl225w (light purple) and NAC (Egd1: light green, Egd2: dark green). Model on the right was rotated ∼50° and highlights the interaction of the ubiquitin-associated (UBA) domain of Egd2 (outlined in gold) with Ypl225w. (B) Extracts derived from the untagged strain (*EGD2*, *YPL225W*) or those tagged at the indicated endogenous loci (*EGD2-13xMyc*, *YPL225W-3xFLAG*) were subjected to FLAG IP. Elutions were resolved by SDS-PAGE followed by immunoblotting (IB) with the indicated antibodies. (C) The indicated proteins were prepared and subject to FLAG IP as described in the STAR Methods. Total and eluted (IP) fractions were analyzed by SDS-PAGE and Coomassie staining or immunoblotting (IB) for the Myc tag on Egd2. Ypl225w-3xFLAG was quantitatively captured under these conditions. Dashes indicate cropping from the same gel to remove irrelevant lanes. NAC refers to recombinant Myc-Egd2•10xHis-Egd1. (D) FLAG IP of the indicated proteins was carried out as in panel C. Dashes indicate cropping from the same Coomassie-stained gel to remove irrelevant lanes. (E) As in Fig. 2a but with the indicated yeast extracts. (F) As in Fig. 2b but with *EGD2* or *egd2*Δ extracts supplemented with the indicated concentrations of recombinant NAC proteins. Dashes indicate cropping from the same gel to remove irrelevant lanes. (G) As in panel F but with *egd2*Δ extracts supplemented with the indicated concentrations of Ypl225w-3xFLAG.

First, we observed that Ypl225w-3xFLAG co-immunoprecipitated a C-terminally Myc-tagged version of Egd2 (Fig. 4b). Next, we attempted to pull down recombinant NAC with recombinant Ypl225w-3xFLAG using anti-FLAG magnetic beads but found no evidence of a stable interaction (Fig. 4c). We observed a similar negative result when we incubated Ypl225w-3xFLAG with salt-washed ribosomes purified from yeast cell extracts, which notably lacked stoichiometric amounts of NAC (Fig. 4c). Strikingly, Ypl225w-3xFLAG co-immunoprecipitated near quantitative amounts of both NAC and ribosomes following incubation of all components together (Fig. 4c). This interaction was abolished when we replaced wild-type NAC (NAC^WT^) with a NAC mutant lacking the UBA domain (NAC*^uba^*^Δ^) altogether or containing point mutations within the UBA domain (NAC^L165A^ and NAC^V166A^) that our ColabFold model predicted would disrupt the interaction with Ypl225w (Fig. 4d and Extended Data Fig. 4a).

To examine the requirement for NAC during eEF1A biogenesis in our cell-free system, we generated *egd2*Δ cell extracts and found evidence of severe eEF1A misfolding relative to wild-type extracts but no discernible effect on Sup35 folding (Fig. 4e and Extended Data Fig. 4b). Importantly, co-translational add-back of recombinant NAC^WT^ but not NAC*^uba^*^Δ^ or NAC^L165A^ ^+ V166A^ mutants restored folding to normal levels (Fig. 4f). Next, we reasoned that if NAC facilitates timely co-translational capture of eEF1A nascent chains, adding supraphysiological concentrations of Ypl225w-3xFLAG should bypass the need for NAC *in vitro*. Consistently, titration of *egd2*Δ cell extracts with additional recombinant Ypl225w-3xFLAG progressively suppressed their eEF1A folding defect (Fig. 4g). Lastly, we found that Ypl225w-3xFLAG cross-linking to V160 RNCs was diminished in *egd2*Δ extracts in a manner that was rescued by add-back of NAC^WT^ but not NAC*^uba^*^Δ^ (Extended Data Fig. 4c). In sum, these findings strongly argue that NAC facilitates co-translational eEF1A folding via an SRP-like tethering mechanism directed at Ypl225w.

### Ypl225w mediates eEF1A folding using an N-terminal α-helix with a hydrophobic patch

To address how Ypl225w facilitates eEF1A folding, we examined Ypl225w’s long NaH, which is predicted to insert itself into the switch helical region of eEF1A’s DI (Fig. 1b). We focused on a hydrophobic patch on NaH comprising several highly conserved residues, which we rendered more hydrophilic by mutating phenylalanine at position 19 to alanine (F19A; Fig. 5a and Extended Data Fig. 5a). We found multiple lines of evidence that F19A ablates Ypl225w’s chaperone function. First, replacing wild-type Ypl225w-3xFLAG at the endogenous locus with the stably-expressed F19A mutant allele resulted in a *ypl225w*Δ-like HSR induction (Fig. 5b and Extended Data Fig. 5b). Second, we analyzed Ypl225w^F19A^-3xFLAG interactors via SDS-PAGE, Western blotting, and negative stain electron microscopy and found a strong reduction in Ypl225w-bound eEF1A, ribosomal proteins, and ribosomal particles (Fig. 5c and Extended Data Fig. 5c). Third, co-translational addback of Ypl225w^F19A^-3xFLAG to *ypl225w*Δ extracts, even at supraphysiological concentrations, failed to restore trypsin resistance to eEF1A (Fig. 5d and Extended Data Fig. 5d). Fourth, Ypl225w^F19A^-3xFLAG yielded a much weaker interaction with V160 RNCs by our cross-linking assay (Fig. 5e). Lastly, we examined the effect of F19A under fully-defined, *in vitro* conditions and observed that Ypl225w-3xFLAG quantitatively pulls down recombinant eEF1A DI, as well as full-length eEF1A, whereas Ypl225w^F19A^-3xFLAG does not interact with either (Fig. 5f). As further evidence of pull-down specificity, we observed no stable interaction between Ypl225w-3xFLAG and Sup35 DI (Extended Data Fig. 5e). Taken together, these data demonstrate that NaH mediates Ypl225w’s interaction with its folding client using a selective, hydrophobic interface.

**Fig. 5.**
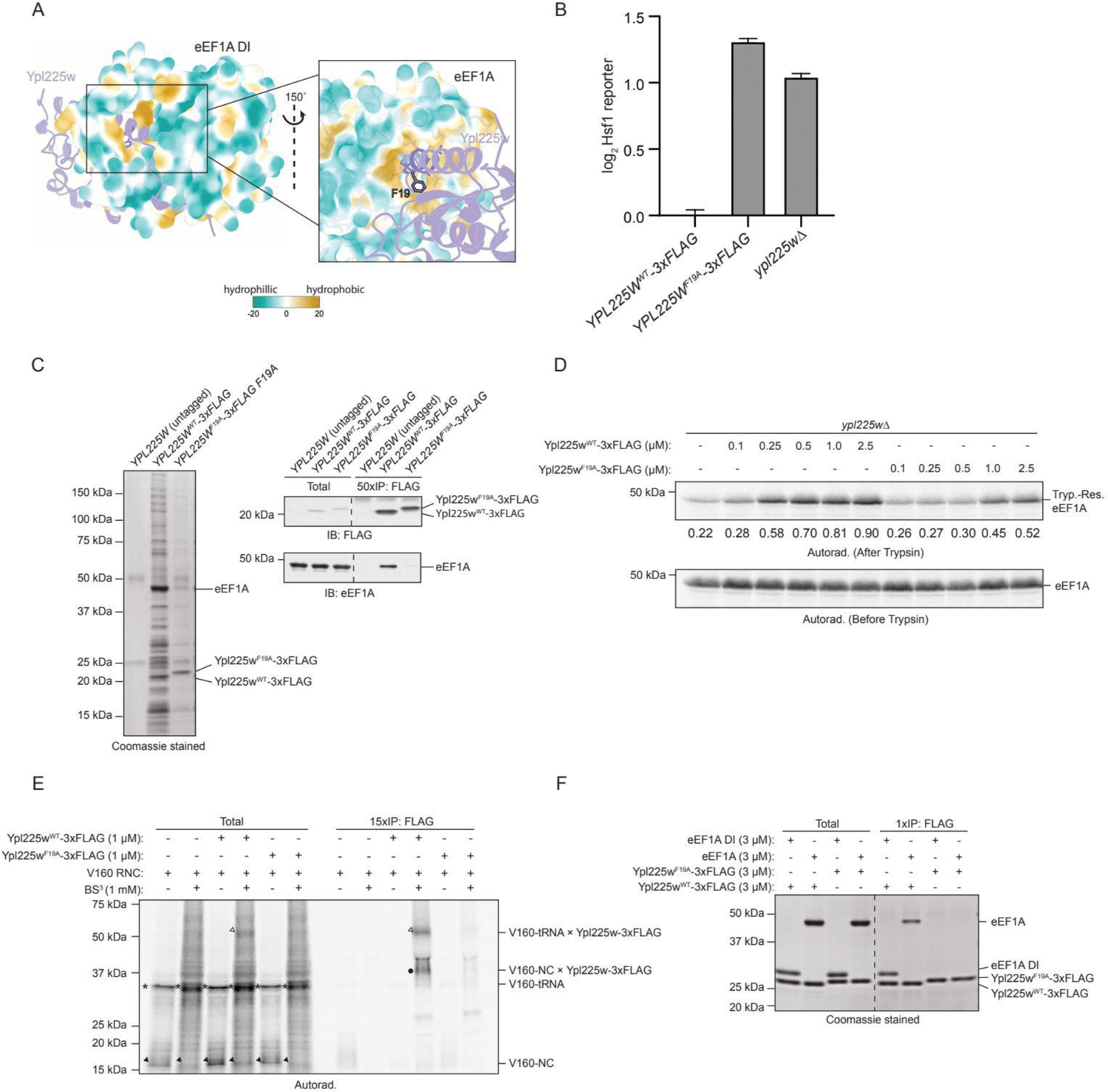
Ypl225w uses its hydrophobic N-terminal α-helix to facilitate eEF1A folding. (A) ColabFold models of eEF1A DI in surface representation bound to Ypl225w (light purple ribbon). Hydrophobicity was defined using ChimeraX molecular lipophilicity potential (“mlp”) command with default settings. Inset on the right was rotated ∼150° and illustrates the hydrophobic interaction of the eEF1A DI SwI region with the NaH of Ypl225w. The F19 residue on the helix is highlighted in black. (B) Indicated strains carrying the Hsf1 reporter (*4xHSE-YFP*) were analyzed by flow cytometry as in Fig. 1c and normalized to *YPL225W^WT^-3xFLAG* values. (C) Untagged (*YPL225W*), *YPL225W^WT^-3xFLAG*, and *YPL225W^F19A^-3xFLAG* samples were analyzed as in Fig. 4b but with Coomassie staining (left), as well as immunoblotting (IB, right) with the indicated antibodies. (D) As in Fig. 2b but with *ypl225w*Δ extracts supplemented with the indicated concentrations of Ypl225w^WT^-3xFLAG or Ypl225w^F19A^-3xFLAG proteins. (E) As in Fig. 3d but with *ypl225w*Δ extracts supplemented with Ypl225w^WT^-3xFLAG or Ypl225w^F19A^-3xFLAG proteins and cross-linked where indicated (lanes with BS^3^). Filled arrow: V160-NC, asterisk: V160-tRNA, filled circle: V160-NC x Ypl225w-3xFLAG, open arrow: V160-tRNA x Ypl225w-3xFLAG. (F) FLAG IP of the indicated proteins as in Fig. 4c. eEF1A indicates the full-length protein. Dashes indicate cropping from the same Coomassie-stained gel to remove irrelevant lanes.

### Ypl225w is a GTP-dependent eEF1A foldase

Ypl225w is predicted to interact with DI switch regions in their nucleotide-free state raising the intriguing possibility that GTP binding and perhaps even GTP hydrolysis could drive release of Ypl225w from eEF1A DI. To test this idea, we first established that GTP indeed disrupts endogenous binding of Ypl225w-3xFLAG to eEF1A by adding excess GTP during the isolation procedure (Extended Data Fig. 6a). Additionally, recombinant Ypl225w-3xFLAG was impaired in its ability to pull down eEF1A DI in the presence of GTP (Extended Data Fig. 6b). To confirm that this effect is driven by formation of a GTP-bound eEF1A conformation, we used a GTP-binding defective version of DI (DI^D156N^)^31,32^. As expected, Ypl225w-3xFLAG interaction with DI^D156N^ was resistant to disruption by GTP (Extended Data Fig. 6b). Next, we developed a bead-based assay for monitoring the kinetics of GTP-driven release of DI-immobilized Ypl225w-3xFLAG. We pre-formed protein complexes in the absence of nucleotides and bound them to anti-FLAG magnetic beads. Following GTP addition, we observed fast release of DI^WT^ which occurred in a concentration-dependent manner (Fig. 6a-d). The addition of GDP and non-hydrolyzable GTP analogs, but not GMP or the guanosine base alone also stimulated release of DI^WT^, suggesting that the negatively charged β-phosphate group of GTP is critical for driving release but that GTP hydrolysis is not required (Extended Data Fig. 6c). To additionally probe the base specificity of these nucleotide effects, we monitored the effect of xanthosine triphosphate (XTP), the deaminated derivative of GTP, on DI release. In earlier work, eEF1A^WT^ was shown to have little affinity for XTP relative to GTP, whereas eEF1A^D156N^ had the opposite specificity for these two nucleotides^31,32^. As expected, GTP but not XTP caused rapid release of DI^WT^ from Ypl225w-3xFLAG, whereas the reverse was true for DI^D156N^ (Fig. 6c-d).

**Fig. 6.**
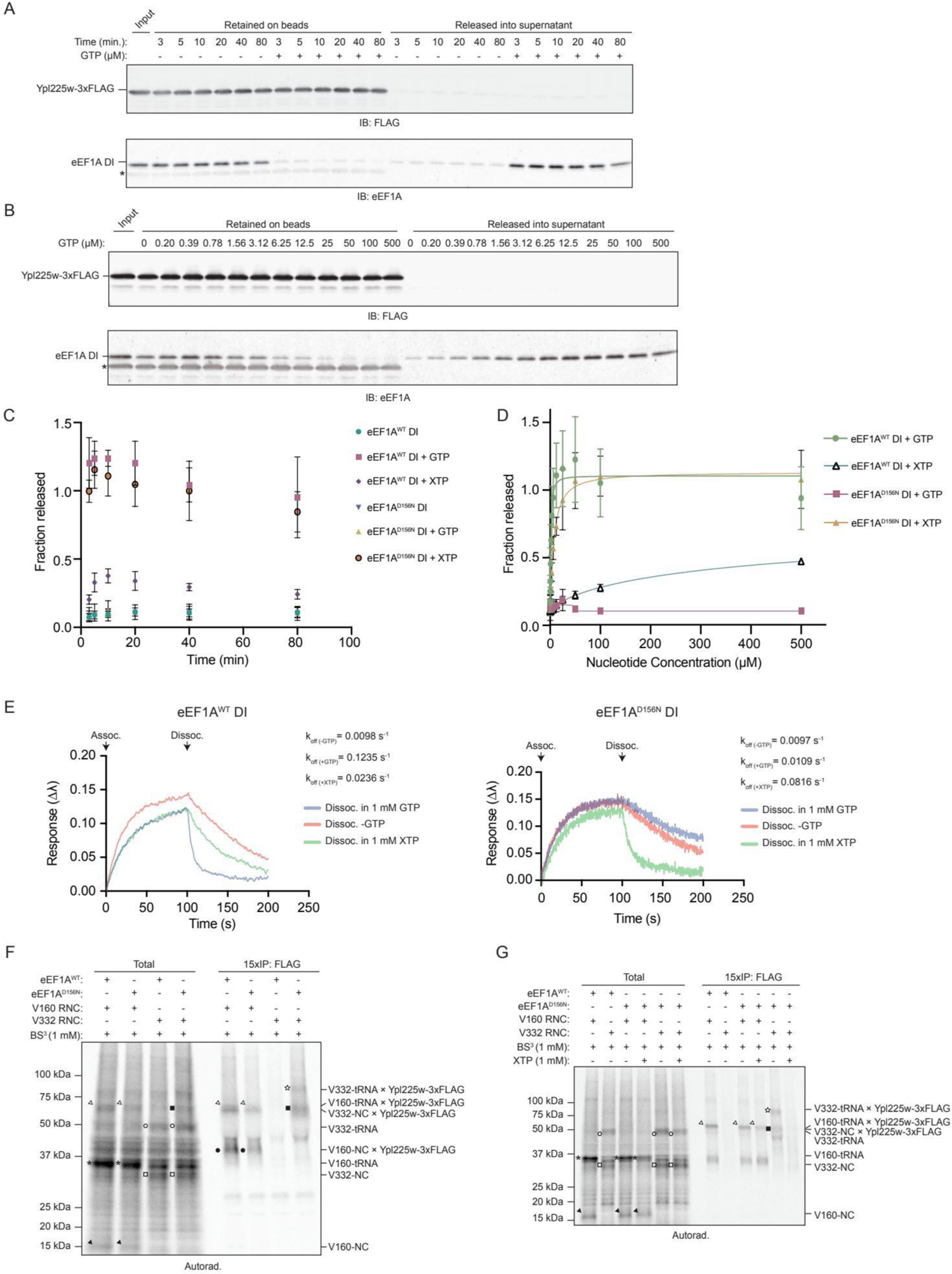
Ypl225w release is predicated on eEF1A DI binding to GTP. (A) eEF1A DI•Ypl225w-3xFLAG complexes were first immobilized on beads (∼300 nM Ypl225w and ∼150 nM eEF1A DI), which were subsequently incubated with or without 100 µM GTP. At the indicated time points, samples were subjected to magnetic separation and removal of the supernatant fraction, followed by elution of the beads with FLAG peptide. Samples were resolved by SDS-PAGE and visualized by immunoblotting (IB). The ‘Input’ lane represents beads that were directly eluted with FLAG peptide after the initial immobilization of eEF1A DI•Ypl225w-3xFLAG. ‘Fraction released’ is the eEF1A DI signal in each supernatant over the input signal. * Asterisk indicates FLAG bleed-through signal in anti-eEF1A immunoblot. (B) As in panel A but beads were incubated with the indicated concentration of GTP for 10 minutes. (C) Line plot showing quantification of DI^WT^ or DI^D156N^ released from Ypl225w-3xFLAG beads as in panel A but in the presence or absence of 100 µM GTP or XTP for the indicated times. Data represents mean ± standard deviation of three replicates. (D) Line plot showing quantification of DI^WT^ or DI^D156N^ released from Ypl225w-3xFLAG beads at different concentrations of nucleotide as in panel B but with either GTP or XTP. Data represents mean ± standard deviation of three replicates. A sigmoidal (4PL) curve was fitted to the data in GraphPad Prism. (E) The association (‘Assoc.’) of DI^WT^ or DI^D156N^ to Ypl225w-3xFLAG-coated BLI tips and subsequent dissociation (‘Dissoc.’) in 1 mM GTP, 1 mM XTP, or buffer (‘-GTP’) was measured in triplicate and plotted as shown. The dissociation rate constants (k_off_) for the different reactions are indicated on the plot. (F) As in Fig. 3d but with either eEF1A^WT^ or eEF1A^D156N^ mRNA templates. Filled arrow: V160-NC, asterisk: V160-tRNA, filled circle: V160-NC x Ypl225w-3xFLAG, open arrow: V160-tRNA x Ypl225w-3xFLAG, open square: V332-NC, open circle: V332-tRNA, filled square: V332-NC x Ypl225w-3xFLAG, open star: V332-tRNA x Ypl225w-3xFLAG. (G) As in panel F but 3 minutes prior to cross-linking, reactions were supplemented with 1 mM XTP where indicated.

The bead-release data above favor a chaperone “foldase” model in which Ypl225w primes DI for GTP binding as opposed to a competing view in which Ypl225w is a “holdase” that releases nucleotide-free DI to undergo a subsequent folding event coupled to GTP binding. To exclude the possibility – which would be consistent the above holdase view – that DI is rapidly dissociating and re-binding in the above assays with bead-immobilized Ypl225w-3xFLAG such that GTP actually blocks re-binding, we used bio-layer interferometry (BLI), a technique for measuring the kinetics of molecular interactions in real time^33^. First, we immobilized Ypl225w-3xFLAG on BLI protein G tips that we loaded previously with anti-FLAG antibody. Next, we monitored DI^WT^ and DI^D156N^ binding to the tips before shifting them into a complex dissociation regime where we could directly examine the kinetic effects of GTP or XTP. In further support of the foldase model, GTP induced almost instantaneous dissociation of DI^WT^, while XTP was only marginally faster than spontaneous dissociation (Fig. 6e). In the case of DI^D156N^, we observed the anticipated reversal of nucleotide kinetic effects with XTP now leading to almost instantaneous release of DI^D156N^ while GTP was ineffective (Fig. 6e). These data establish that GTP actively dissociates Ypl225w from its client rather than working passively in a client “trapping” mechanism.

Lastly, we exploited eEF1A^D156N^’s preference for XTP over GTP in the more physiologically-relevant, cell-free system to test two related predictions of the foldase model. First, reduced binding of GTP to growing D156N RNCs should cause them to stay persistently bound to Ypl225w, representing a chaperone recycling defect. Second, this defect should be resolved in the presence of XTP. We validated the first prediction by repeating our cross-linking assay using D156N V160-RNC and V332-RNCs. In contrast to the wild-type control in which only the V160-RNC interacted with Ypl225w-3xFLAG, we observed strong cross-links to both D156N V160- and V332-RNCs (Fig. 6f). To test the second prediction, we included XTP either during D156N RNC translation or following inhibition of further protein synthesis by cycloheximide. In both instances, we observed the expected outcome in which XTP selectively disrupted Ypl225w’s interaction with V332-RNCs (Fig. 6g and Extended Data Fig. 6d). These two pieces of data fit well with our working model that Ypl225w stabilizes eEF1A DI nascent chains in a near-native conformation primed for GTP-driven folding.

## Discussion

Co-translational protein folding in prokaryotes depends on Trigger Factor (TF), an ATP-independent chaperone unique to prokaryotes^34^. TF is also the only known chaperone in prokaryotes that binds ribosomes even though other ATPase chaperones, including Hsp70, can associate with nascent chains^34^. By contrast, studies of co-translational protein folding in eukaryotes have shown that RAC exerts control over the Hsp70 ATPase cycle by a complex and still poorly understood mechanism^5,6,10,35^. In addition, eukaryotic ribosomes are bound to stoichiometric amounts of NAC^3^. The precise role of NAC in folding has remained unclear but recent work has clearly illustrated NAC’s versatility in mediating other kinds of protein biogenesis decisions, ranging from ER targeting to enzymatic processing of nascent N-termini^3,8,9^. In this study, we provide evidence for a mechanistic model (Fig. 7), in which the UBA domain of NAC facilitates ribosomal recruitment of Ypl225w, a previously uncharacterized ATP-independent chaperone, to mediate co-translational folding of eEF1A, one of the most abundant proteins in eukaryotes.

**Fig. 7.**
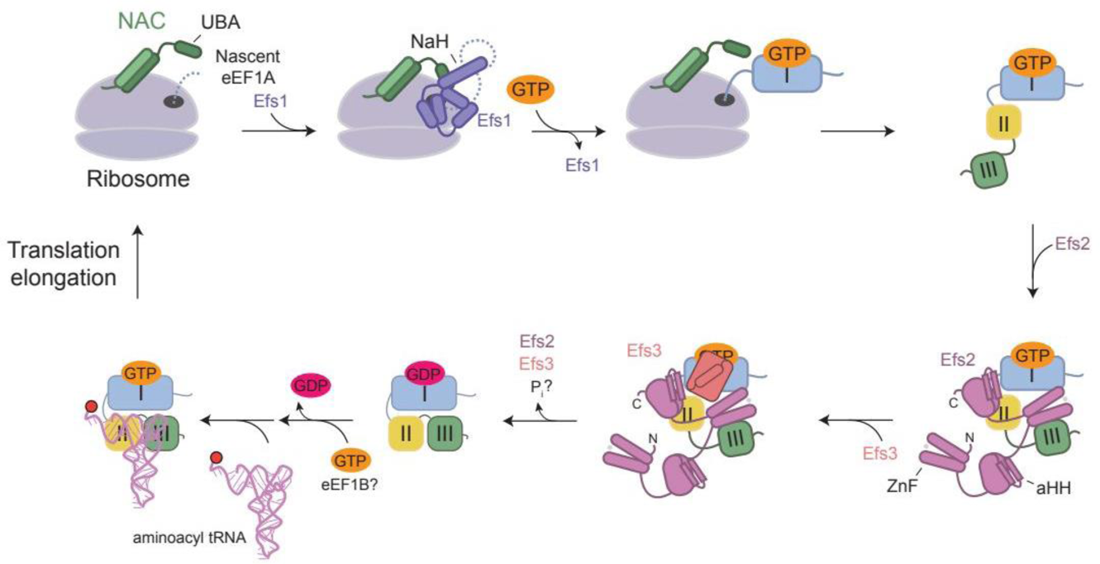
Model for the GTP-dependent chaperone cycle of eEF1A biogenesis. Cartoon model emphasizing the GTP-driven chaperone mechanisms of the eEF1A Folding Sequence (EFS) en route to formation of eEF1A ternary complexes competent for translation elongation. See Discussion for further details.

Our model offers the first mechanistic explanation for NAC’s long-suspected role in protein folding but resembles the mechanism by which metazoan NAC directs recruitment of SRP to nascent ER signal sequences^9^. Interestingly, an interaction between the *C. elegans* orthologs of Egd2 (NACB) and Ypl225w (Pbdc1) has been previously reported^36^, thus arguing that we have uncovered a broadly conserved chaperone system. Further supporting this view are previously reported proteomic interactions between the human orthologs of eEF1A (EEF1A) and Ypl225w (PBDC1)^36–39^, as well as our own cross-species complementation data (Extended Data Fig. 1g). An important future goal will be to examine the extent to which NACB UBA mutants defective specifically for Pbdc1 binding disrupt proteostasis in metazoans.

As part of our model, we favor the simplest view that Ypl225w only becomes stably associated with the ribosome following emergence of Ypl225w’s minimal client binding site, which is predicted to reside within the switch regions in the N-terminal half of eEF1A’s DI. Our data also suggest that the kinetics of this client recognition event are hastened by Ypl225w’s additional interactions with NAC UBA and an undefined ribosome binding site. Ypl225w then maintains the nascent chain in a state primed for GTP binding upon emergence of additional DI structural elements needed to form a bona fide GTP-binding pocket. Finally, elongation enables a switch-like mechanism in which GTP-binding induces DI folding concomitant with Ypl225w release from the ribosome. It is conceivable that GTP can drive this last step without requiring a full-fledged DI, similar to the proposed mechanism by which ATP drives vectorial folding of CFTR’s nucleotide-binding domain 1^40^. We also cannot exclude more elaborate models in which NAC first senses short eEF1A nascent chains – as hopeful Ypl225w clients – before helping Ypl225w dock to the ribosome. Cryo-EM analysis of purified Ypl225w-bound RNCs, as well as approaches for selective ribosome profiling in vivo^41^ should help resolve these issues in the future.

Our work also sets the stage for further exploring the evolutionary origins behind eEF1A’s dependence on a dedicated, co-translational chaperone system. Specifically, we have found computational and experimental evidence that the highly-related DI of Sup35 folds independently of Ypl225w. Thus, one future strategy would be to screen for eEF1A DI mutant sequences that fold along a smoother, Sup35-like folding pathway. Such chaperone bypass might have naturally taken place during evolution in plant species such as *Arabidopsis thaliana*, an organism lacking discernible Ypl225w homologs. While speculative, this intriguing possibility is consistent with our folding simulations of nascent *A. thaliana* eEF1A showing native-like intermediates reminiscent of Sup35 rather than the non-native structure of yeast eEF1A (Extended Data Fig. 7a-b).

Lastly, the addition of Ypl225w to the repertoire of eEF1A-specific chaperones motivated us to define the eEF1A Folding Sequence (EFS) as a series of sequential, GTP-driven steps mediated by three ATP-independent chaperones (re-named Efs1-3 according to their order of action; Fig. 7). Besides reflecting the major role of these proteins in eEF1A folding, we hope this new nomenclature will also correct the long-standing but erroneous conflation of the Ypl225w/PBDC1 Pfam family (PFO4669) with unrelated plant-specific enzymes involved in polysaccharide biosynthesis.

## Acknowledgements

We thank Jeffrey Prince, Kate Guerin, and Oliver Rancu for their help with the initial characterization of Ypl225w. We would like to thank Mei Chen and Steven Kowalski of the Harvard Center for Mass Spectrometry for their help with mass spectrometry analysis and preparation. We also would like to thank the Harvard Center for Macromolecular Interactions for their help with bio-layer interferometry. This work was supported by the National Institutes of Health (R35-GM127136 to V.D.) and the National Science Foundation Graduate Research Fellowships Program (DGE1745303 to I.M.S.).

## Author Contributions

Conceptualization, I.M.S. and V.D.; methodology, I.M.S., D.R., K.P., A.J.M., A.R.N., and C.T.; formal analysis, I.M.S., D.R., K.P., A.R.N.; investigation, I.M.S., D.R., K.P., A.J.M., and A.R.N.; resources, A.W.M. and V.D.; writing – original draft, I.M.S., D.R., and V.D.; writing – reviewing and editing, all authors; supervision, V.D., E.I.S., A.W.M., and V.D.; funding acquisition, I.M.S. and V.D.

## Ethics declarations

The authors declare no competing interests.

## Methods

### Yeast Strains

All strains used in this study are listed in Supplementary Table 2. Cells were grown or cultured at 30°C unless otherwise indicated.

### Plasmids

All plasmids used in this study are listed in Supplementary Table 3.

### Oligos

All oligos used in this study are listed in Supplementary Table 4.

### Antibodies

For immunoblotting, primary antibodies were used at the indicated dilution and fluorescent secondary antibodies were used at a 1:10,000 dilution. Primary antibodies include the following: rabbit-anti-eEF1A ([1:10,000 - IB], ED7001, Kerafast Inc., Boston, MA), mouse-anti-FLAG M2 ([1:2000-1:5000 - IB], F3165, MilliporeSigma, Burlington, MA), mouse-anti-Pgk1 ([1:2000 - IB], 459250, Thermo Fisher Scientific, Waltham, MA), mouse-anti-Myc 9E10 ([1:10,000 - IB], 13-2500, Thermo Fisher Scientific, Waltham, MA), and rabbit-anti-Hexokinase ([1:2000 - IB], U.S. Biological Life Sciences, Swampscott, MA).

### Immunoblotting

Samples were analyzed via SDS-PAGE, transferred to a nitrocellulose membrane, and blocked using 5% skim milk in TBST (50 mM Tris-HCl pH 7.4, 150 mM NaCl, 0.1% Tween 20, 0.25 mM EDTA) for 1 hour. Primary antibodies at the indicated dilutions (see Antibodies section) were used to probe membranes overnight at 4°C. Prior to secondary antibody incubation (see Antibodies section), membranes were washed three times using TBST (10 minutes per wash). Secondary antibodies were visualized using a LI-COR Odyssey scanner.

### Autoradiography

After separation via SDS-PAGE, gels were soaked in a fixing solution consisting of 50% methanol and 10% acetic acid for 30 minutes before being rinsed in a drying solution consisting of 30% methanol and 5% glycerol for 30 minutes. Gels were then dried under a vacuum for 2 hours. Next, gels were exposed using phosphor-screens (GE Healthcare) overnight and subsequently scanned on a Sapphire Biomolecular Imager (Azure Biosciences).

### Microscopy

#### eEF1A-GFP aggregation assay

VDY6176 and VDY6206 cells were grown to saturation overnight in synthetic complete (SC) media containing 2% glucose. After back-dilution to an OD_600_ of 0.1 in SC + 2% glucose, cells were grown for 3-4 hours until they reached mid-log phase. Cells were then imaged using a Nikon TI inverted microscope with a 100x oil-immersion objective (1.45 NA), a Yokogawa dual spinning disk confocal unit, and a Hamamatsu ImagEM EM-CCD camera with a 488 nm laser (GFP). Images were captured with MetaMorph. Maximum intensity z-projections were used to generate the final images from collected Z stacks.

#### eEF1A DI-GFP

VDY6205 cells expressing *pGAL1*-eEF1A DI-GFP (pVD2659) were grown overnight in SC + 2% raffinose. After back-dilution to an OD_600_ = 0.1 in SC + 2% galactose, cells were grown for another 4 hours and imaged as described above.

#### eEF1A DI-GFP/Sup35 DI-GFP

VDY6374 and VDY6375 cells were transformed with pVD3006/3007. Transformants were grown overnight in S-Trp (synthetic media lacking tryptophan) + 2% glucose. Cells were next back-diluted to an OD_600_ of 0.1 and grown for another 4 hours and subsequently imaged as described above.

### Flow cytometry

For flow cytometry, experiments were performed using the FACSymphony A3 analyzer (BD Biosciences) with the 488 nm laser (FITC). 10,000 events were collected for every sample.

### *4xHSE-YFP* reporter of Hsf1 activity

VDY3334 and VDY6165 cells were grown to saturation overnight in SC + 2% glucose. After back-dilution to an OD_600_ of 0.1, cells were grown for 4-5 hours until OD_600_ ∼0.5. Samples were then measured, and medians were collected using Bioconductor packages flowCore and flowViz.

To determine the HSR of Ypl225w^F19A^-3xFLAG-expressing cells relative to Ypl225w^WT^-3xFLAG-expressing cells and *ypl225w*Δ cells, VDY6292, VDY6334, and VDY6341 cells were grown to saturation overnight in SC + 2% glucose. Subsequent dilution, growth, measurement, and analysis was performed as described above.

#### eEF1A-GFP levels

VDY6176 and VDY6206 cells were grown to saturation in SC + 2% glucose media. Cells were back-diluted to an OD_600_ of 0.1 in SC + 2% glucose and grown until OD_600_ = 0.5. Samples were immediately measured. 10,000 cells were measured for each sample and medians were collected as described above.

#### eEF1A DI-GFP levels

VDY6205 and VDY6240 cells were grown overnight in SC + 2% raffinose media. Cells were back-diluted to an OD_600_ = 0.1 in SC + 2% galactose and grown for 4 hours. Samples were measured and sample medians were analyzed as described above.

#### GCN4 uORFs-GFP reporter

VDY6374 and VDY6375 were transformed with pVD2746. Transformants were grown overnight in S-Ura (synthetic media lacking uracil) + 2% glucose and back-diluted to an OD_600_ of 0.2. Cells were then grown for 4-5 hours until mid-log phase was reached, and samples were subsequently measured. Median collection and analysis was performed as described above.

### Growth assays

#### Spot assays

VDY6285, VDY6287, VDY6335, and VDY6336 cells were grown to mid-log phase and back-diluted to an OD600 of 0.4. The first spot of each row represents the undiluted culture and subsequent spots were taken from serial dilutions (1:10).

#### Growth curves

VDY3334 and VDY6165 cells were grown to mid-log phase before back-dilution to an OD600 of 0.05 in 200 µL of YPD rich media in a 96-well plate. Cells were grown with shaking for 24 hours at 30°C with OD_600_ measurements collected at 10 minute-intervals. Three replicates were measured for each sample.

### Polysome profiling

#### Ypl225w-3xFLAG EDTA treatment

VDY6217 cells were grown logarithmically in YPD before back-dilution to reach an OD_600_ of 0.5. Next, cells were harvested by rapid vacuum filtration and snap freezing in liquid nitrogen. The frozen paste was mixed with frozen polysome lysis buffer (20 mM Tris pH 8.0, 70 mM KCl, 5 mM MgCl_2_, 100 µg/mL cycloheximide, 1 mM DTT, 1% Triton X-100, 0.025 U/µL Turbo DNase [AM2238, Thermo Fisher Scientific]) with 1 mL lysis buffer added per 500 OD_600_ units. The frozen mix was then cryogenically lysed using the Retsch MM 400 mixer mill and the resultant grindate was thawed for 2 minutes at 30°C with intermittent flicking and placed on ice. 40 mM EDTA was added during this thawing step for samples that received this treatment. Samples were clarified via centrifugation for 10 minutes at 16,000 x g and 4°C and the RNA concentration was normalized across samples. 1 mg of RNA was loaded onto linear 10-50% sucrose gradients and centrifuged in a Beckmann Coulter SW-41 rotor for 3 hours at 30,000 RPM and 4°C. Gradients were fractionated using a BioComp Gradient Station and the absorbance at 260 nm was measured during fractionation. Trichloroacetic acid (TCA) was added to fractionated samples to a final concentration of 20% and samples were stored at −20°C overnight. The next day, samples were spun down for 30 minutes at 4°C, washed three times with ice-cold acetone (where each wash consisted of a 10-minute 4°C centrifugation step), and vacuum-dried. The protein pellet was solubilized in SDS loading buffer and separated via SDS-PAGE before immunoblotting with the indicated antibodies.

#### Ypl225w-3xFLAG + Tef-Off

VDY6238 cells were grown logarithmically in YP-Gal (2% galactose) media before being divided 1:1 with either fresh YP-Gal media or YPD (and supplemented to 2% glucose final). Galactose- and glucose-treated cells were then grown for 3 hours to a final OD_600_ of 0.5 before being harvested, fractionated, and analyzed as described above.

### Immunoprecipitations

#### Ypl225w-3xFLAG from cell lysates

Cells containing endogenously tagged Ypl225w^WT^-3xFLAG or Ypl225w^F19A^-3xFLAG were grown in 1.5 L cultures to OD_600_ of 1.8 in liquid YPD media. Cells were spun down at 3500 x g for 15 minutes at 4°C, washed in 50 mL double-distilled water twice, and resuspended in 1 mL (per gram of pellet) resuspension buffer (1.2% PVP-40, 20 mM HEPES pH 7.4, 1x protease inhibitor cocktail [Roche], 1% solution P [2 mg Pepstatin A, 90 mg PMSF, 5 mL 100% ethanol], 1 mM DTT). Cells were spun down at 3500 x g for 15 minutes to remove all traces of buffer. The cell paste was placed into a syringe and was pushed out to be frozen as “noodles” in a 50 mL conical tube filled with liquid nitrogen. Liquid nitrogen was decanted from the tube and frozen “noodles” were stored at −80°C until cryogenically lysed using a Retsch PM100 ball mill. Next, 250 mg of powder was thawed at 4°C followed by resuspension in 1 mL of HIP buffer (40 mM HEPES-KOH pH 7.5, 110 mM KOAc, 2 mM MgCl_2_, 1% Triton X-100, 0.1% Tween, 1x protease inhibitor cocktail [Roche], 1% solution P, 1 mM DTT). The lysate was next passed through a Whatman 25 mm GD/X Disposable filter (Cat No. 6888-2527) and equal volumes of the filtered lysates were added to 62.5 μL Protein G Dynabeads pre-bound to 3.75 μL of anti-FLAG M2 antibody. Samples were subsequently agitated for 30 minutes at 4°C. Beads were washed twice with 750 µL of HIP buffer. After the final wash, protein was eluted from beads with 40 μL of 1 mg/mL 3xFLAG peptide in HIP buffer at room temperature for 30 minutes. Samples were analyzed via SDS-PAGE followed by immunoblotting with the indicated antibodies.

For Extended Data Fig. 6, 1 mM GTP and the regeneration system (1 mM ATP, 10 mM creatine phosphate, 0.04 mg/mL creatine kinase) were supplemented as indicated to all HIP wash buffer steps.

#### Purified Ypl225w-3xFLAG with eEF1A, eEF1A DI, or Sup35 DI

Pure eEF1A, eEF1A DI, or Sup35 DI was diluted in Buffer A (20 mM HEPES pH 7.4, 100 mM KoAc, 2 mM Mg(OAc)_2_, 2 mM DTT) with 5% glycerol at 2x the final concentration. Ypl225w-3xFLAG and nucleotide (if present) was added for 30 minutes so that the final concentration of each protein was 3 µM and the nucleotide concentration was 100 µM or 1 mM. An aliquot was removed as the total. Samples were added to Protein G Dynabeads pre-conjugated to anti-FLAG antibody (∼1 µL antibody per 16.6 µL beads) resuspended in 1 reaction volume of Buffer A with 5% glycerol. Samples were agitated at 4°C for 30 minutes and the unbound material was removed, and samples were washed 3x in 1 mL cold wash buffer (Buffer A with 5% glycerol and 0.05% Tween). Samples were eluted after agitating for 30 minutes at RT with 1 mg/mL 3xFLAG peptide in Buffer A with 5% glycerol. Equal amounts of totals and IPs were added to 2x SDS/Urea sample buffer (20 mM Tris-HCl pH 6.8, 4 M Urea, 2.5% SDS, 0.05 mM EDTA, 1% Beta-mercaptoethanol, 10% glycerol), heated at 80°C, and analyzed by SDS-PAGE and Coomassie staining.

#### Purified Ypl225w-3xFLAG with ribosomes and NAC

A final concentration of 125 A_260_ units of high salt-washed ribosomes, 7.5 µM NAC or NAC mutant protein, and 3 µM Ypl225w-3xFLAG protein were incubated as indicated together for 30 minutes at room temperature. An aliquot was removed as the total. Samples were added to Protein G Dynabeads pre-conjugated to anti-FLAG antibody (∼1 µL antibody per 16.6 µL beads) resuspended in 1 reaction volume of Buffer A with 5% glycerol. Samples were agitated at 4°C for 30 minutes, unbound material was removed, and samples were washed 3x in 1 mL cold wash buffer (Buffer A with 5% glycerol and 0.05% Tween). Samples were eluted after agitating for 30 minutes at room temperature with 1 mg/mL 3xFLAG peptide in Buffer A with 5% glycerol. Equal amounts of totals and IPs were added to 2x SDS/Urea sample buffer and analyzed by SDS-PAGE and Coomassie staining or immunoblotting with the indicated antibodies.

#### eEF1A DI release assays

For release assays to monitor eEF1A DI dissociation from Ypl225w-3xFLAG, the following changes were made: reactions were set up so that the final concentration of eEF1A and Ypl225w would be ∼150 nM and 300 nM respectively in 24 µL after splitting the washed beads into 13 tubes. One sample was set aside, and the remaining 12 were incubated in 24 µL Buffer A + 5% glycerol with the indicated added components for the indicated times at room temperature with agitation. The supernatant was removed, and the beads for all 13 samples were eluted with 24 µL 1 mg/mL 3xFLAG peptide in Buffer A + 5% glycerol as before. Equal amounts of supernatant and eluate from the beads were added to 2x SDS/Urea sample buffer and analyzed by SDS-PAGE and immunoblotting.

### Yeast *in vitro* transcription and translation

#### Generation of capped mRNA

The mMESSAGE mMACHINE T7 kit (Invitrogen) was used as described previously to generate capped mRNAs from PCR templates^42^. PCR products encoding *TEF2* were generated from pVD2609 using primers oVD13102 and oVD13103. Nonstop mRNAs were generated from pVD2609 using primers oVD13102 and oVD13432 (V160-RNCs) or oVD13603 (V332-RNCs). D156N versions of eEF1A V160/V332 nonstop mRNAs were generated using pVD2860 as a template. PCR products encoding *SUP35* were generated from genomic *S. cerevisiae* DNA.

#### Preparation of cell-free translation extracts

Yeast *in vitro* translation (IVT) extracts were prepared as described previously^12^. Briefly, 1.5 L YPD media of indicated strains were grown to an OD_600_ ∼ 1.8-2.0. Cells were centrifuged at 3,000 RPM for 15 minutes in a JS-4.2 rotor at 4°C. Next, pellets were first resuspended in DEPC-treated water, centrifuged, and then resuspended in Buffer A (30 mM HEPES pH 7.4, 100 mM KoAc, 2 mM Mg(OAc)_2_) with 2 mM DTT. Finally, 1 mL Buffer A with 2 mM DTT, 14% glycerol (Buffer ADG), and 2x cOmplete protease inhibitor cocktail (PIC) tablet (Roche, Basel, Switzerland) was added per 6 grams of cell weight and frozen via a ‘popcorn’ method (frozen drop by drop in liquid nitrogen) before being lysed in a Retsch PM100 ball mill. The powder was thawed at 4°C and centrifuged at 13,336 x g for 10 minutes at 4°C. The resultant supernatant was transferred to an SW-55 rotor and centrifuged at 49,000 RPM for 30 minutes at 4°C with no brake. The middle two layers were removed for subsequent desalting (using five connected Hi-Trap desalting columns (GE Healthcare, Chicago, IL, USA) on an AKTA Pure FPLC system). Fractions with A_260_ > 60 were pooled, RNase-treated (and subsequently deactivated using EGTA), and snap frozen.

#### eEF1A trypsin protection time course

Each single IVT reaction consisted of 7.5 µL of nuclease-treated extracts, 2 µL Buffer ADG, 2.5 µL of 6x translation buffer (132 mM HEPES-KOH, pH 7.4, 720 mM KOAc, 9 mM Mg(OAc)_2_, 4.5 mM ATP, 0.6 mM GTP, 150 mM creatine phosphate [Roche], 0.24 mM of each amino acid but lacking methionine [Promega, Madison, WI], 10.2 mM DTT), 0.5 µL creatine kinase [20 mg/mL in 50% glycerol, Roche], 0.5 µL RiboGuard [Lucigen, Middleton, WI, USA], 1 µL ^35^S-Methionine [PerkinElmer, Waltham, MA, USA], and 1 µL of 200 ng/µL capped mRNA. Briefly, IVT reactions (once translation had initiated) were incubated for 45 minutes at room temperature (unless otherwise stated). Cycloheximide was added to 100 µg/mL to arrest translation and an aliquot was taken for ‘Before Trypsin’ gel samples. The remainder of the reaction was treated with 50 µg/mL trypsin (Promega) and incubated at room temperature for the indicated times. Samples were quenched using an equal volume of 2x SDS/Urea sample buffer before being heated for 3 minutes at 70°C. Samples were analyzed via SDS-PAGE, autoradiography, and immunoblotting.

#### Generation of RNCs, cross-linking, and denaturing IPs

For the generation of RNCs, four IVT reactions were used per sample and each sample was programmed with nonstop eEF1A mRNA encoding a truncation of either 160 or 332 amino acids. Translation reactions were incubated for 15 minutes at room temperature prior to arrest with cycloheximide. Next, samples were then treated with either water or BS^3^ to 1 mM final concentration. Cross-linking samples were incubated on ice for 90 minutes in the dark. After 90 minutes, 20 mM Tris pH 6.8 was added to quench the cross-linking reagent. SDS and DTT were then added to each sample to 1% SDS and 0.5 mM DTT final concentration to denature non-covalent interactions, and samples were heated subsequently for 10 min at 65°C. Next, cold IP buffer (buffer ADG + 0.1% Triton X-100) was added to dilute denaturant 1:10 (e.g., 450 µL IP buffer added to 50 µL IVT reaction) and samples were added to protein G Dynabeads pre-conjugated to anti-FLAG antibody (0.5 µL antibody per 10 µL beads). Samples were immunoprecipitated by gentle nutation for 30 minutes at 4°C and Dynabeads were then washed twice in cold IP buffer. Samples were eluted using 70°C-heated 1x SDS sample buffer (40 mM Tris-HCl pH 6.8, 5% SDS, 0.1 mM EDTA, 1% Beta-mercaptoethanol, 0.1 mg/mL bromophenol blue, 20% glycerol). Samples were analyzed via SDS-PAGE, autoradiography, and immunoblotting.

### Recombinant protein expression and purification

#### eEF1A DI and Sup35 DI

eEF1A DI or Sup35 DI constructs were cloned into pET29b vectors with a Myc-6xHis C-terminal tag. Rosetta (DE3) cells were transformed with these vectors and grown at 37°C in Terrific Broth (TB) media until OD_600_ of 0.5. The cultures were moved to 4°C for 1 hour and then supplemented with 0.4 mM IPTG (US Biological Life Sciences) to induce protein expression at 16°C for 16 hours. Cells were harvested at 3,500 x g for 20 minutes and resuspended in PBS. After a second spin, cell pellets were frozen in liquid nitrogen and stored at −80°C until purification.

Cell pellets were thawed at 4°C and resuspended in lysis buffer (50 mM Tris pH 7.5, 150 mM NaCl, 2% glycerol, 2mM MgCl_2_) supplemented with 2 mM beta-mercaptoethanol, protease inhibitor cocktail [Roche], and 1 mM PMSF. Cells were lysed using a sonicator set to 40% power for 5x 30 second intervals on ice with 1 minute on ice in between. Lysed cells were spun down at 20,000 x g for 15 minutes at 4°C in a JLA 16.25 rotor. Supernatants were loaded on 1 mL equilibrated HisPur Ni-NTA resin (ThermoFisher) in gravity flow columns twice. The resin was washed sequentially with 20 mL wash buffer 1 (50 mM Tris pH 7.5, 500 mM NaCl, 2% glycerol, 2 mM MgCl_2_, 2 mM beta-mercaptoethanol), 20 mL wash buffer 2 (50 mM Tris pH 7.5, 150 mM NaCl, 2% glycerol, 2 mM MgCl_2_, 10 mM imidazole, 2 mM beta-mercaptoethanol), and 20 mL wash buffer 3 (50 mM Tris pH 7.5, 150 mM NaCl, 2% glycerol, 2 mM MgCl_2_, 50 mM imidazole, 2 mM beta-mercaptoethanol). The resin was then incubated with 3 mL of elution buffer (50 mM Tris pH 7.5, 150 mM NaCl, 2% glycerol, 2 mM MgCl_2_, 250 mM imidazole, 2 mM beta-mercaptoethanol) for 5 minutes before eluting.

Eluate was concentrated to <1 mL and applied to an equilibrated Superdex 200 10/300 column for gel filtration in SEC buffer (40 mM HEPES pH 7.5, 110 mM KOAc, 2 mM MgCl_2_, 2% glycerol, 0.5 mM TCEP). Peak fractions were aliquoted, frozen in liquid nitrogen, and stored at −80°C.

#### Ypl225w-3xFLAG

Ypl225w-3xFLAG constructs were cloned into pET29b vectors with a 3xFLAG-3c-6xHis C-terminal tag. Rosetta (DE3) cells were transformed with these vectors and grown at 37°C in Terrific Broth (TB) media until OD_600_ of 0.5. The cultures were then supplemented with 1 mM IPTG (US Biological Life Sciences) to induce protein expression at 37°C for 4 hours. Cells were harvested at 3,500 x g for 20 minutes and resuspended in PBS. After a second spin, cell pellets were frozen in liquid nitrogen and stored at −80°C until purification.

Cell pellets were thawed at 4°C, resuspended in B1 Lysis Buffer (20 mM sodium phosphate pH 8.0, 300 mM NaCl, 20 mM imidazole) supplemented with benzonase (Millipore Sigma), 5 mM beta-mercaptoethanol, protease inhibitor cocktail [Roche], and 0.2 mM PMSF. Cells were lysed with 5 passes on an Emulsiflex cell disruptor at ∼12-15k PSI. Lysed cells were spun down at 28,000 x g for 30 minutes at 4°C in a JLA 16.25 rotor. The resultant supernatant was loaded onto an equilibrated 5 mL HisTrap HP column. The column was washed with 10 column volumes of B1 buffer before eluting with a linear gradient of B2 Elution Buffer (20 mM sodium phosphate pH 8.0, 300 mM NaCl, 500 mM imidazole, 5 mM beta-mercaptoethanol) over 5 column volumes. Peak elution fractions were pooled and concentrated to 2.5 mL with a 10 kDa cutoff filter before desalting with a PD10 desalting column into B5 SEC Buffer (50 mM Tris pH 8.0, 200 mM NaCl, 10% glycerol, 0.5 mM TCEP). 1 mL of desalted sample was applied to an equilibrated Superdex 200 10/300 column for gel filtration in B5 SEC buffer. Peak fractions were aliquoted, frozen in liquid nitrogen, and stored at −80°C.

#### NAC

NAC constructs were cloned into pETDuet vectors with N-terminal Myc-Egd2 and 10xHis-Egd1 tags. Rosetta (DE3) cells were transformed with these vectors and grown at 37°C in Luria (LB) media until OD_600_ of 0.6. The cultures were then supplemented with 1 mM IPTG (US Biological Life Sciences) to induce protein expression at 37°C for 3 hours. Cells were harvested at 3,500 x g for 20 minutes and resuspended in IMAC-A buffer (50 mM HEPES pH 7.5, 300 mM NaCl, 20 mM imidazole, 1 mM DTT, 5% glycerol) supplemented with 1 mM PMSF. After a second spin, cell pellets were frozen in liquid nitrogen and stored at −80°C until purification.

Cell pellets were thawed at 4°C and resuspended in IMAC-A Buffer supplemented with 1 mM PMSF. Next, cells were lysed using a sonicator set to 40% power for 3x 30 second intervals on ice. Lysed cells were spun down at 40,000 x g for 30 minutes at 4°C in a JA 25.5 rotor. Resultant supernatants were loaded on 1 mL equilibrated HisPur Ni-NTA resin (ThermoFisher) in gravity flow columns and washed with 15 mL of IMAC-A buffer. The protein was eluted in 3 mL IMAC-B buffer (50 mM HEPES pH 7.5, 300 mM NaCl, 500 mM imidazole, 1 mM DTT, 5% glycerol). The eluate was concentrated to 0.5 mL with a 10 kDa cutoff filter before being applied to an equilibrated Superdex 200 10/300 column for gel filtration in SEC buffer (50 mM HEPES pH 7.5, 150 mM NaCl, 0.5 mM TCEP). Peak fractions were combined, concentrated, aliquoted, frozen in liquid nitrogen, and stored at −80°C.

### Purification of native eEF1A from yeast

Native eEF1A was purified from *S. cerevisiae* as previously described^12^. VDY465 (WT) cells were grown in large cultures to an OD_600_ of 1.8 in liquid YPD media. 3 L of cells were spun down at 3500 x g for 20 minutes at 4°C, washed in 50 mL double-distilled water twice, and resuspended in 1 mL per gram of pellet of resuspension buffer (1.2% PVP-40, 20 mM HEPES pH 7.4, 1x protease inhibitor cocktail [Roche], 1% solution P [2 mg Pepstatin A, 90 mg PMSF, 5 mL 100% ethanol], 1 mM DTT). Cells were spun down twice at 3500 x g for 10 minutes to remove all buffer. The resultant cell paste was placed into a syringe and was pushed out to be frozen as “noodles” in a 50 mL conical tube filled with liquid nitrogen. Liquid nitrogen was decanted from the tube and frozen “noodles” were stored at −80°C until cryogenically lysed using a Retsch PM100 ball mill.

Next, 8 g of frozen grindate was thawed in 80 mL I-100 buffer (20 mM Tris pH 7.5, 0.1 mM EDTA, 100 mM KCl, 25% glycerol, 1 mM DTT) supplemented with 1x protease inhibitor cocktail (Roche). Lysate was spun at 8,000 x g for 5 minutes at 4°C in a JA 25.5 rotor. Supernatant was transferred into new tubes and spun at 20,000 x g for 15 minutes at 4°C in JA 25.5 rotor. The supernatant was then spun at 54,400 RPM for 106 minutes at 4°C in Ti-70 ultracentrifuge rotor. Next, the supernatant was incubated with 10 g of pre-equilibrated DEAE resin (in buffer I-100) for 1 hour while agitated on a tilt table at 4°C. The unbound fraction was removed by centrifugation. The resin was washed with an additional 20 mL I-100 buffer, spun down, and the wash was added to the unbound fraction. The unbound fraction and wash were incubated with 60 mL pre-equilibrated CM-sepharose slurry (in buffer I-100) for 1 hour while agitated on a tilt table at 4°C. A Buchner funnel with Whatman type 1 paper (∼11 µm pore size) was used to separate the unbound fraction. The resin was washed with 40 mL I-100 before removing from the Buchner funnel, resuspended in 60 mL I-100 with solid KCl supplemented to 500 mM final concentration of KCl, and incubated for 1 hour while agitated on a tilt table at 4°C. Eluate was collected using the Buchner funnel, and resin was washed with an additional 20 mL buffer I-100 supplemented with solid KCl for a final concentration of 500 mM KCl, which was combined with the eluate. Next, the eluate was concentrated with a 30 kDa cutoff concentrator to ∼5 mL, filtered with a 0.22 µm filter and desalted into I-50 buffer (20 mM Tris pH 7.5, 0.1 mM EDTA, 50 mM KCl, 25% glycerol, 1 mM DTT) using 4 x 5 mL sequential HiTrap desalting columns. ∼8 mL of flow-through was collected and applied onto a Source 15S 4.6/100 column. The column was washed with 40 column volumes (CVs) of I-50 buffer before eluting with a linear gradient of I-300 buffer (20 mM Tris pH 7.5, 0.1 mM EDTA, 300 mM KCl, 25% glycerol, 1 mM DTT) over 20 CVs. Peak fractions containing eEF1A were pooled and concentrated to ∼2 mL with a 30 kDa cutoff concentrator. The sample was next applied onto a HiPrep 16/60 Sephacryl S-100 column. Native eEF1A eluted as the second peak. Fractions were pooled and concentrated with a 30 kDa cutoff concentrator. Aliquots were frozen in liquid nitrogen and stored at −80°C.

### Purification of high-salt washed ribosomes

VDY6292 cells were grown in large cultures to an OD_600_ 1.8 in liquid YPD media. 4.5 L of cells were spun down at 3500 x g for 20 minutes at 4°C, washed in 50 mL double-distilled water twice, and resuspended in 1 mL per gram of pellet of resuspension buffer. Cells were spun down twice at 3500 x g for 10 minutes to remove all buffer. Cell paste was placed into a syringe and was pushed out to be frozen as “noodles” in a 50 mL conical tube filled with liquid nitrogen. Liquid nitrogen was decanted from the tube and frozen “noodles” were stored at −80°C until cryogenically lysed using a Retsch PM100 ball mill.

Next, powder was thawed in 25 mL of ribosome lysis buffer (1x Buffer A, 1 mg/mL heparin sodium salt (Sigma)) supplemented with 2 mM DTT, 1x protease inhibitor cocktail (Roche), and 0.1 mM PMSF at 4°C. The lysate was cleared by centrifugation at 20,000 x g for 30 minutes at 4°C in a JA 25.5 rotor. Next, the supernatant was brought up to 35 mL with ribosome lysis buffer supplemented with 2 mM DTT, 1x protease inhibitor cocktail (Roche), and 0.1 mM PMSF, and ∼17.5 mL of the solution was layered onto two 3 mL low-salt sucrose cushions (1x Buffer A, 100 mM KCl, 500 mM sucrose) before centrifugation for 106 minutes at 60,000 x rpm in a Ti-70 rotor at 4°C. The supernatant was removed carefully, and each pellet was resuspended carefully in 1.5 mL high salt wash buffer (1x Buffer A, 600 mM KCl, 1 mg/mL heparin, 2 mM DTT).The volumes were combined, brought up to 9 mL in a 50 mL falcon tube and left overnight at 4°C to fully resuspend. 3 x 3 mL sample was layered onto 3x 250 µL high-salt sucrose cushions (1x Buffer A, 600 mM KCl, 500 mM sucrose), before centrifugation at 100,000 x rpm for 30 minutes at 4°C. Pellets were rinsed once in freezing buffer (1x Buffer A, 10% glycerol, 2 mM DTT) before resuspending in 100 µL freezing buffer. Aliquots were frozen in liquid nitrogen and stored at −80°C.

### Bio-layer interferometry

Biolayer interferometry was performed on an Octet RED384 instrument (Sartorius). Anti-FLAG M2 was immobilized on protein G biosensors (Sartorius Cat. No. 18-5082) at 5 µg/mL (in binding buffer: Buffer A with 5% glycerol, 2 mM DTT, and 0.05% Tween-20) for 120 seconds, after which a baseline measurement was performed in a new well with the same buffer for 30 seconds. Sensors were then moved into 1 µM Ypl225w-FLAG for 30 seconds to capture Ypl225w-FLAG. After Ypl225w-FLAG binding, sensors were moved into binding buffer with 1 mM GTP (Roche), 1 mM XTP (TriLink Biotechnologies), or no nucleotide to acquire baseline measurement for 30 seconds. Then sensors were moved into 300 nM eEF1A DI (DI^WT^ or DI^D156N^) domain in binding buffer without nucleotide for the association phase. After 100 seconds the sensors were moved back into baseline wells with or without nucleotides to measure eEF1A DI domain dissociation in the presence of different nucleotides. Probes were regenerated in 0.1 mM Glycine pH 1.5 and neutralized in binding buffer before rebinding anti-FLAG M2 and reusing for subsequent measurements. Measurements were obtained in quadruplicate or triplicate for each condition. Data were analyzed in the included Octet RED384 data analysis software. Briefly, each sensor was subjected to double reference subtraction. First, a reference sensor with no Ypl225w-3xFLAG was subtracted from each sensor. Then, a reference well with no eEF1A DI domain was subtracted from sensors subjected to eEF1A DI domain binding. The data were corrected to adjust signal to the same value between the association and dissociation steps to correct for optical artifacts caused by moving sensors between wells.

### Negative stain of YPL225W, YPL225W^WT^-3xFLAG, and YPL225W^F19A^-3xFLAG strains

Samples were stained using a 1.5% (w/v) uranyl formate solution and imaged on a ThermoScientific Tecnai T12 electron microscope equipped with a LaB6 filament, operated at 120 kV and at 59000x magnification. The microscope is equipped with a Gatan UltraScan 895 (4k x 4k) CCD camera.

### ColabFold modeling of Ypl225w•eEF1A and NAC•Ypl225w•eEF1A complexes

ColabFold version 1.5.2 was used with 5 models, 20 seeds, 20 recycles and the default parameters with msa_mode: “MMseqs2 (UniRef+Environmental)”, and model_type: “AlphaFold2-multimer-v3” to model the *S. cerevisiae* eEF1A•Ypl225w and eEF1A DI•Ypl225w complexes. For the *S. cerevisiae* eEF1A DI•Ypl225w•Egd2•Egd1 complex, ColabFold version 1.5.2 was run with the same parameters but with 1 seed. Sequences were obtained from Uniprot. eEF1A DI contains residues 1-239. Figure 1B highlighted residues on eEF1A are: 36-75 (Switch I), 95-110 (Switch II), and 14-21 (P-loop). For Figure S4, the UBA-resembling domain of Ef-Ts was shown as residues 1-40. Models are provided in Supplementary Item 1. Consurf was used to determine conservation scores for Ypl225w residues and CLUSTAL Omega was used for sequence alignment as shown in Figure S5A. Molecular graphics and analyses for all models were performed with UCSF ChimeraX.

### Ypl225w-3xFLAG immunoprecipitation mass spectrometry and analysis

#### Sample preparation

The samples were prepared using filter aided sample preparation (FASP). Protein samples were loaded onto the 10k filter and centrifuged at 13,500 x g for 10 minutes and followed by a wash with 100 mM triethylammonium bicarbonate (TEAB). Proteins were reduced by 20 mM tris(2-carboxyethyl)phosphine (TCEP) for 1 hour at 37 °C followed by centrifugation as described above, and then alkylated in 40 mM iodoacetamide for 1 hour in darkness at the room temperature followed by centrifugation as described above. Protein digestion was then performed using trypsin with 1:50 ratio (w:w, substrate: enzyme) and incubating overnight at 37 °C. Digested filtrate samples were cleaned up with C18 Spin Tips (Pierce, Thermo Scientific) and reconstituted with 0.1% formic acid prior to LC-MS/MS analysis.

#### Liquid chromatography-mass spectrometry method

Samples were analyzed by Lumos Tribrid Orbitrap Mass Spectrometer coupled with Ultimate 3000 nano-HPLC (Thermo Fisher) at the Harvard Center for Mass Spectrometry. Peptides were separated onto a 150 μm inner diameter microcapillary trapping column packed with approximately 2 cm of C18 Reprosil resin (5 μm, 100 Å, Dr. Maisch GmbH, Germany) followed by an analytical column (µPAC, C18 pillar surface, 50 cm bed, Thermo scientific). Separation was achieved by applying a gradient from 7–37% acetonitrile in 0.1% formic acid over 90 minutes at 500 nl/min. Electrospray ionization was enabled by applying a voltage of 2.2 kV using a homemade electrode junction at the end of the microcapillary column and sprayed from metal tips (PepSep, Denmark). The mass spectrometry survey scan was performed in the Orbitrap in the range of 400–1,800 m/z at a resolution of 12×10^4^, followed by the selection of the twenty most intense ions (TOP20) for CID-MS2 fragmentation in the Ion trap using a precursor isolation width window of 2 m/z, AGC setting of 10,000, and a maximum ion accumulation of 100 ms. Singly charged ion species were not subjected to CID fragmentation. The normalized collision energy was set to 35 V and an activation time of 10 ms. Ions in a 10-ppm m/z window around ions selected for MS2 were excluded from further selection for fragmentation for 90 seconds.

#### Analysis of proteomics data

Raw data was then submitted for analysis in Proteome Discoverer 2.5 software (Thermo Scientific). The mass spectrometry data was searched against the Uniprot reviewed *Saccharomyces cerevisiae* database along with known contaminants such as human keratins and common lab contaminants. Quantitative analysis between samples was performed by label-free quantitation (LFQ). Sequest HT searches were performed using the following guidelines: a 10 ppm MS tolerance and 0.6 Da MS/MS tolerance; trypsin digestion with up to two missed cleavages; carbamidomethylation (57.021 Da) on cysteine were set as static modification; oxidation (+15.995 Da) of methionine and protein N-terminus acetylation set as dynamic modification; minimum required peptide length set to ≥ 6 amino acids. At least one unique peptide per protein group is required for identifying proteins. All MS2 spectra assignment FDR of 1% on both protein and peptide level was achieved by applying the target-decoy database search by Percolator.

### Computational screening for interactors of eEF1A

AlphaPulldown in the ‘pulldown’ mode with default parameters was used to screen 3769 yeast proteins conserved in humans (source: DIOPT Ortholog Finder) for candidates forming a binary complex with eEF1A ^16,43^. MSAs were generated using ColabFold/MMSeqs2 prior to running the structure predictions in parallel as a HPC job array. Complexes were filtered for an inter-chain PAE < 5 and a Protein Interaction score based on structural properties of the complex^3^ (PI_score > 0.1)^44^.

### Co-translational folding simulations of eEF1A and Sup35

Markov Chain Monte Carlo (MCMC) simulations with MCPU were conducted to investigate and compare molecular environments during the early stages of protein folding ^19,21^. The initial structures for these simulations were predicted by ColabFold and processed using a minimization process that ran for 50 million steps at a temperature of 0.1 kBT. Subsequently, 600 Monte Carlo (MC) steps were performed, incorporating temperature replica exchange and umbrella sampling, with the temperature varying from 0.4 kBT to 1.0 kBT in increments of 0.025 kBT. Umbrella biasing was also applied with the number of native contacts as a collective variable. For each simulation trajectory, the first 200 million steps were discarded, and the remaining 400 million steps were used for the analysis. Additionally, the Multistate Bennett Acceptance Ratio (MBAR) method was employed to correct for biases introduced during the enhanced sampling, so that the unbiased free energy could be calculated from the simulated trajectories^20^.

### Quantification and Statistical Analysis

#### Flow cytometry analysis

Using BioConductor packages flowCore and flowViz, flow cytometry samples were gated for live, single cells. Samples were normalized to cell size by dividing the raw GFP/YFP intensity (FITC.A) by side scatter (SSC.A). Data was log transformed for histograms and bar graphs.

#### Differential protein abundance analysis of Ypl225w-3xFLAG IP/MS

To detect statistically significant differences in protein abundance, the R package limma (v.3.50.0) was used to calculate the FDR on the Empirical Bayes moderated t-statistic^45^. A protein was defined as significant if FDR < 0.05. The results of this analysis are provided in Supplementary Table 1.

## Data Availability

Further information and requests for resources or reagents should be directed to the lead contact, Vladimir Denic.

---

**Extended Data Fig. 1.**
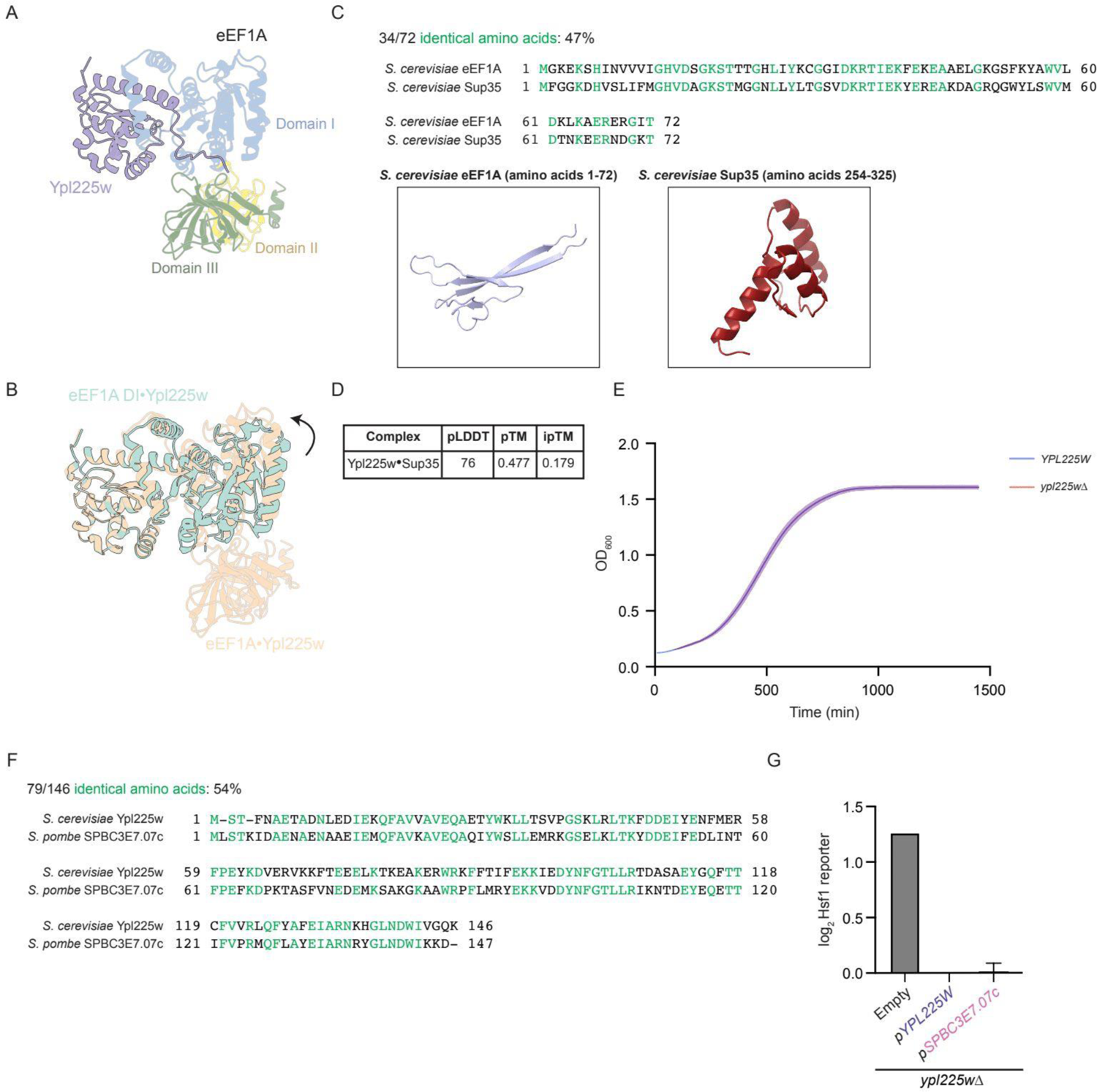
Co-translational modeling simulations of eEF1A reveal a tendency for eEF1A DI to misfold. (A) ColabFold model showing interaction of Ypl225w (light purple) with full-length eEF1A (colored by domain). (B) ColabFold models comparing the predicted full-length eEF1A•Ypl225w (peach) and eEF1A DI•Ypl225w (sea green) complexes aligned on Ypl225w using the “matchmaker” command on ChimeraX. Arrow indicates slight rotation of DI towards Ypl225w in the full-length eEF1A•Ypl225w complex. (C) CLUSTAL Omega multiple sequence alignment of *S. cerevisiae* eEF1A amino acids 1-72 and Sup35 amino acids 254-325 (re-numbered 1-72 with the prion domain (PrD) removed). Identical amino acids are highlighted in green. The residues used for alignment were further used for molecular simulations to compare the folding landscapes of eEF1A and Sup35 (See Methods). Representative snapshots of the minimum free-energy conformation of eEF1A (light blue) and Sup35 (dark red) are shown. (D) ColabFold model confidence scores for Ypl225w•Sup35 complex. (E) Growth curves of WT (*YPL225W*) and *ypl225w*Δ cells inoculated at an initial OD_600_ of 0.05 and grown for 24 hours. The plotted lines represent the average of three biological replicates. The standard deviation is represented as the lighter shading above or below the average line. (F) CLUSTAL Omega multiple sequence alignment of *S. cerevisiae* Ypl225w and *S. pombe* Pbdc1 homolog *SPBC3E7.07c*. Identical amino acids are highlighted in green. (G) *ypl225w*Δ cells carrying the Hsf1 reporter (*4xHSE-YFP*) were transformed with either an empty vector plasmid, a plasmid encoding *S. cerevisiae YPL225W* driven by its endogenous promoter, or a plasmid containing *S. pombe SPBC3E7.07c* driven by *YPL225W*’s promoter. The indicated strains were subsequently measured by flow cytometry and analyzed as in Fig. 1c.

**Extended Data Fig. 2.**
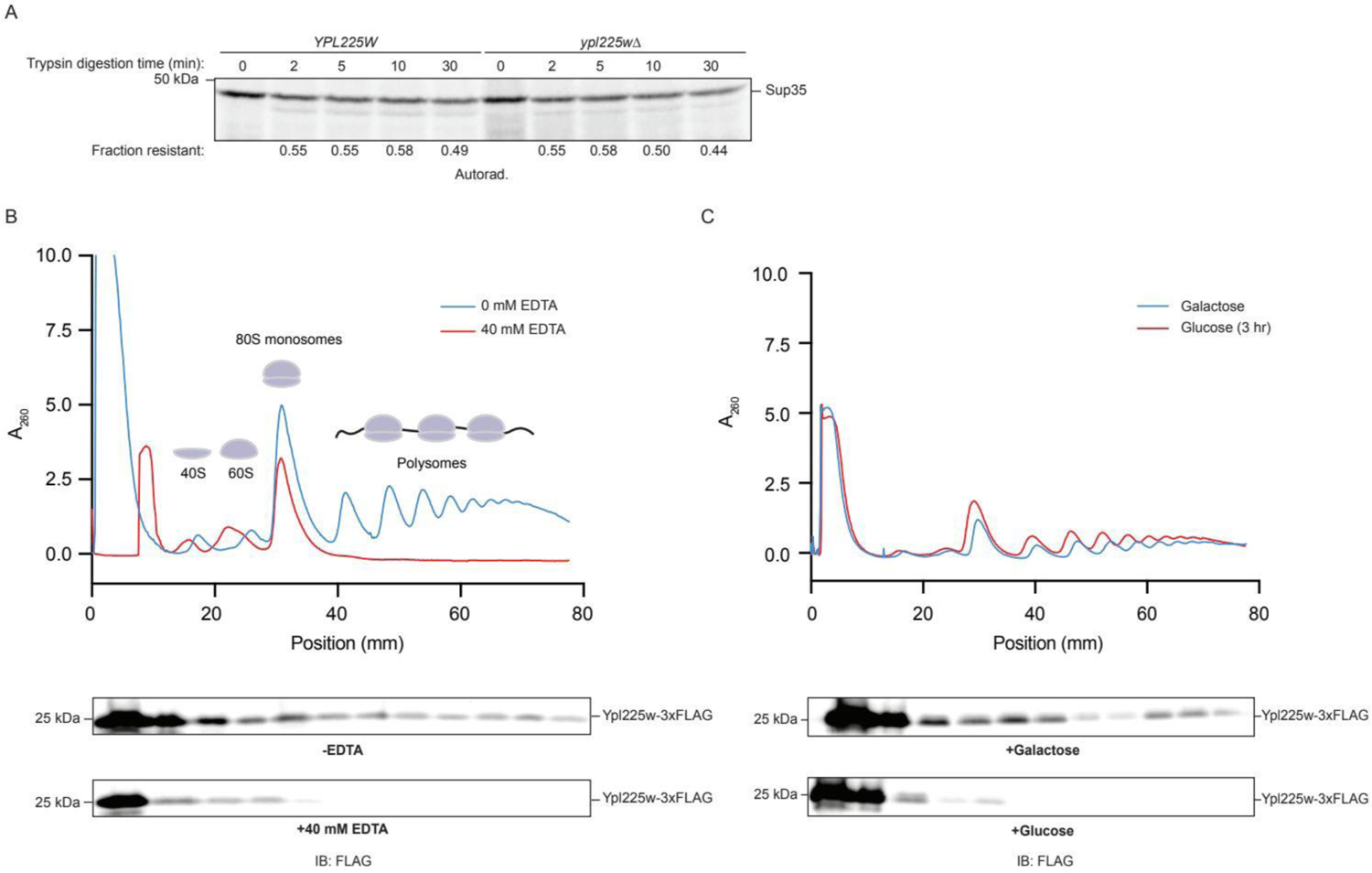
Ypl225w associates with ribosomes in a manner dependent on eEF1A synthesis. (A) As in Fig. 2a but the indicated extracts were programmed with a Sup35 mRNA template. (B) Polysome A_260_ traces of *YPL225W-3xFLAG* cells with (+ 40 mM EDTA) and without (-EDTA) treatment. After sucrose gradient centrifugation and fractionation, fractions were TCA-precipitated, separated by SDS-PAGE, and probed with the indicated antibodies. (C) Polysome A_260_ traces of *YPL225W-3xFLAG* cells with both copies of eEF1A driven by the *GAL1* promoter. Cells were initially grown in YPGal media before being split 1:1 with fresh YPGal (+Galactose) or YPD with additional glucose being added to 2% final (+Glucose). Cells were subsequently grown for another 3 hours before sucrose gradient centrifugation and fractionation. Fractions were TCA-precipitated, separated by SDS-PAGE, and probed with the indicated antibodies.

**Extended Data Fig. 3.**
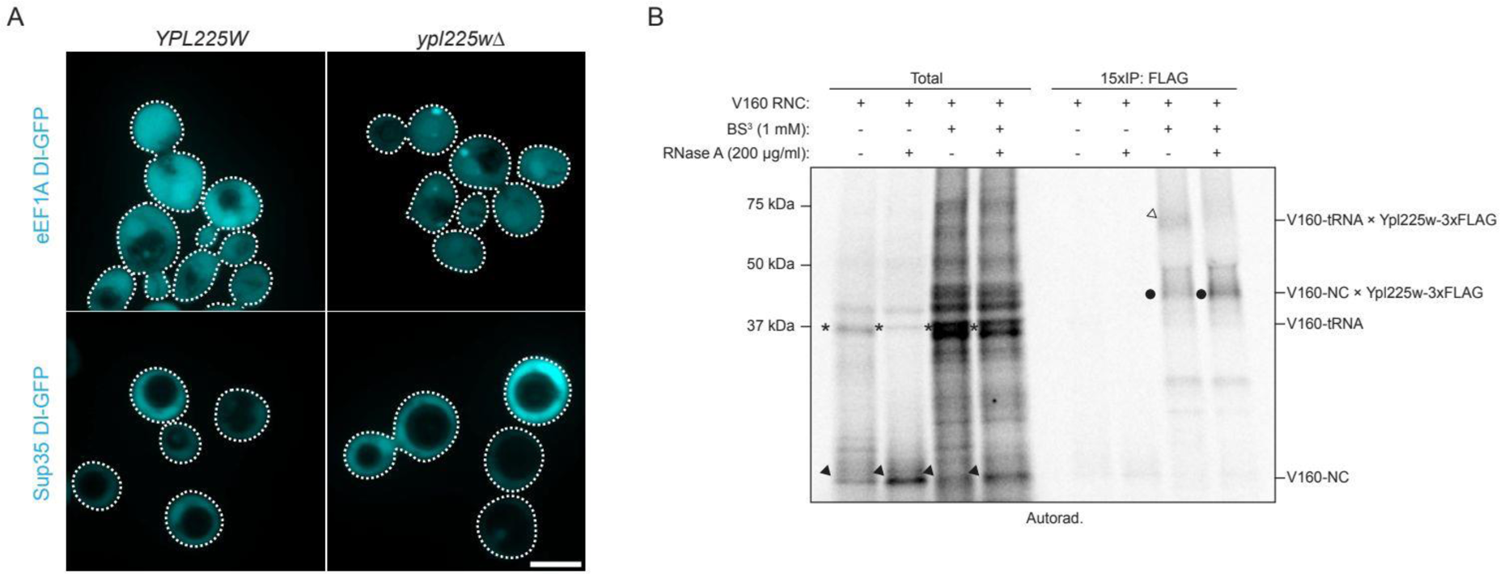
eEF1A DI but not Sup35 DI forms aggregates in cells bearing *ypl225w*Δ. (A) Shown are representative images of WT (*YPL225W*) and *ypl225w*Δ strains containing plasmids expressing eEF1A DI-GFP or Sup35 DI-GFP driven by a *TEF2* promoter. Cells were imaged via confocal microscopy as in Fig. 1d. Scale bar represents 2.5 µm. (B) As in Fig. 3d but extracts were supplemented with RNase A and cross-linked where indicated (lanes with BS^3^). Filled arrow: V160-NC, asterisk: V160-tRNA, filled circle: V160-NC x Ypl225w-3xFLAG, open arrow: V160-tRNA x Ypl225w-3xFLAG.

**Extended Data Fig. 4.**
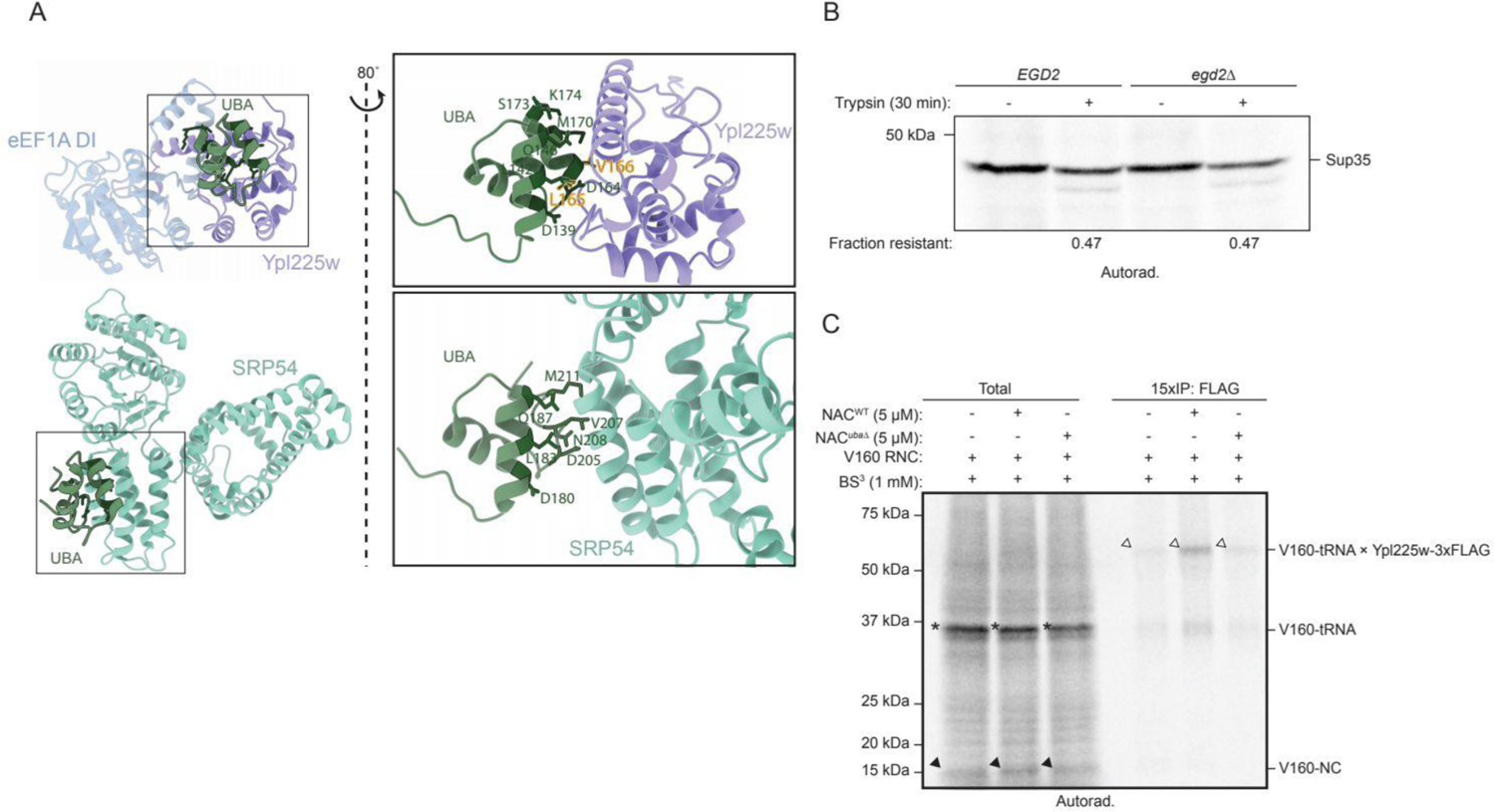
NAC selectively facilitates eEF1A co-translational folding through recruitment of Ypl225w to ribosomes. (A) ColabFold or structural models comparing each UBA interaction interface with insets rotated ∼80° highlighting the UBA interacting residues. Models are aligned on the UBA domain using the ChimeraX “matchmaker” command. The top panels show the ColabFold eEF1A DI (light blue) bound to Ypl225w (light purple) and the UBA domain (dark green) of Egd2 with Egd1 and the majority of Egd2 removed for clarity. UBA residues L165 and V166 are highlighted in gold. The bottom panels show SRP54 (teal) bound to the UBA domain (dark green) of NACA (from pdb 7qwq). (B) As in Fig. 4f, but with *EGD2* or *egd2*Δ extracts translating Sup35 mRNA template. (C) As in Fig. 3d but using *egd2*Δ, *YPL225W-3xFLAG* extracts and supplemented with NAC^WT^ or NAC*^uba^*^Δ^ proteins where indicated. Filled arrow: V160-NC, asterisk: V160-tRNA, filled circle: V160-NC x Ypl225w-3xFLAG, open arrow: V160-tRNA x Ypl225w-3xFLAG.

**Extended Data Fig. 5.**
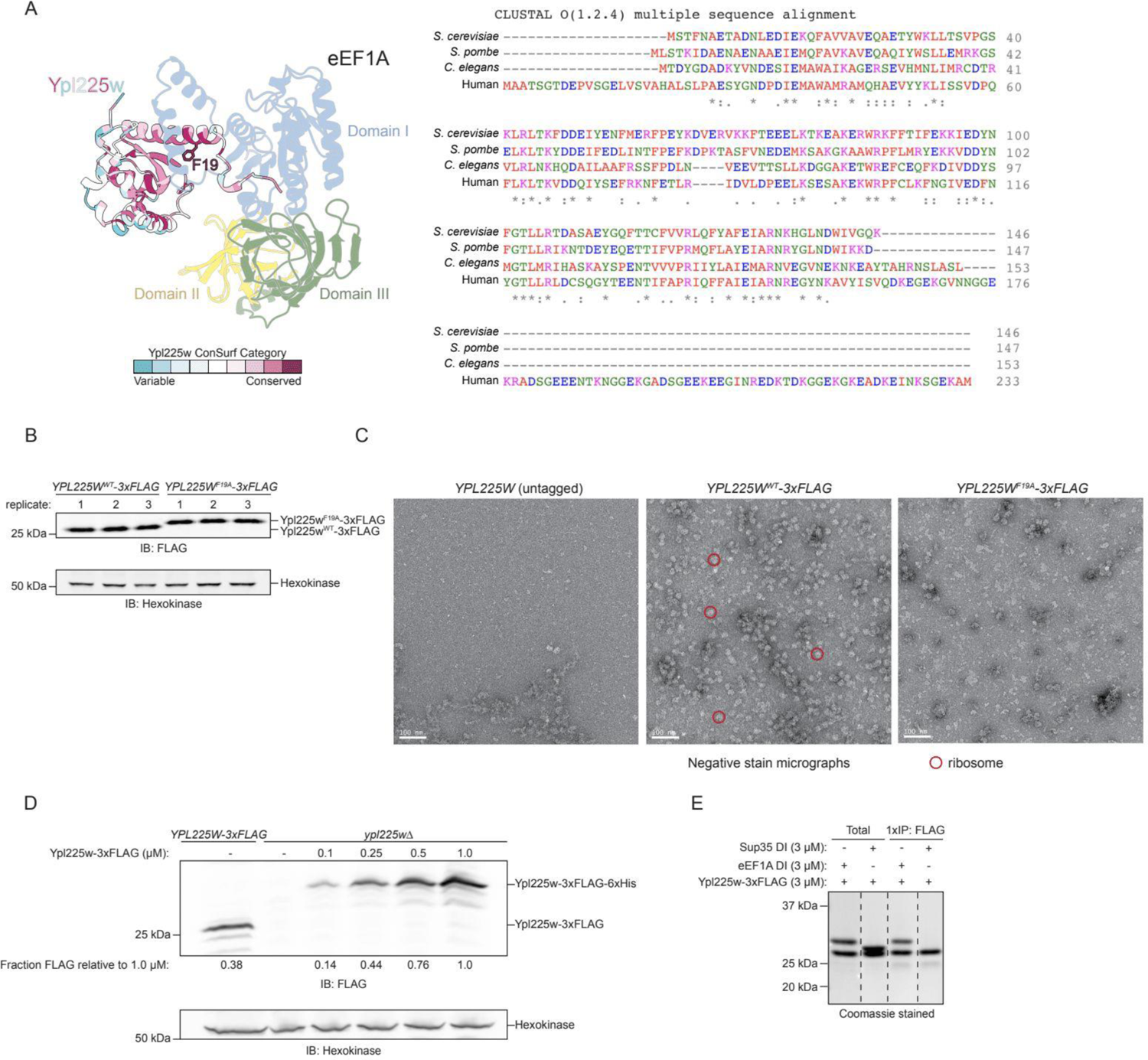
Ypl225w^F19A^ is defective in eEF1A binding and chaperoning. (A) Left: Ribbon diagram of ColabFold model showing eEF1A (colored by domain) bound to Ypl225w F19 residue highlighted (outlined in black). Ypl225w is colored by conservation category as determined by ConSurf, with the color scale indicated. Right: CLUSTAL Omega multiple sequence alignment of *S. cerevisiae* Ypl225w, *S. pombe* Pbdc1 homolog *SPBC3E7.07c*, *C. elegans* Pbdc1, and human Pbdc1 protein sequences. Residues are colored as follows: red (hydrophobic), blue (acidic), magenta (basic), green (hydroxyl, sulfhydryl, amine), gray (unusual). ‘*’ indicates fully conserved, ‘:’ indicates strongly similar, and ‘.’ indicates weakly similar. (B) Samples from Fig. 5b were analyzed by SDS-PAGE and immunoblotting with the indicated antibodies as described in Methods. (C) Representative negative stain EM micrographs of untagged (*YPL225W*), *YPL225W^WT^-3xFLAG*, and *YPL225W^F19A^-3xFLAG* IP samples from Fig. 5c were prepared as described in Methods. Example ribosome particles are indicated with red circles. (D) YPL225W-3xFLAG or *ypl225w*Δ IVT extracts were supplemented with the indicated concentration of recombinant Ypl225w-3xFLAG-6xHis protein and analyzed by SDS-PAGE and immunoblotting for the indicated antibodies. ‘Fraction FLAG relative to 1.0 µM’ is the signal of Ypl225w protein divided by the corresponding hexokinase signal and normalized to the 1.0 µM lane signal. (E) As in Fig. 4c but with the indicated proteins. Dashes indicate cropping from the same Coomassie-stained gel to remove irrelevant lanes.

**Extended Data Fig. 6.**
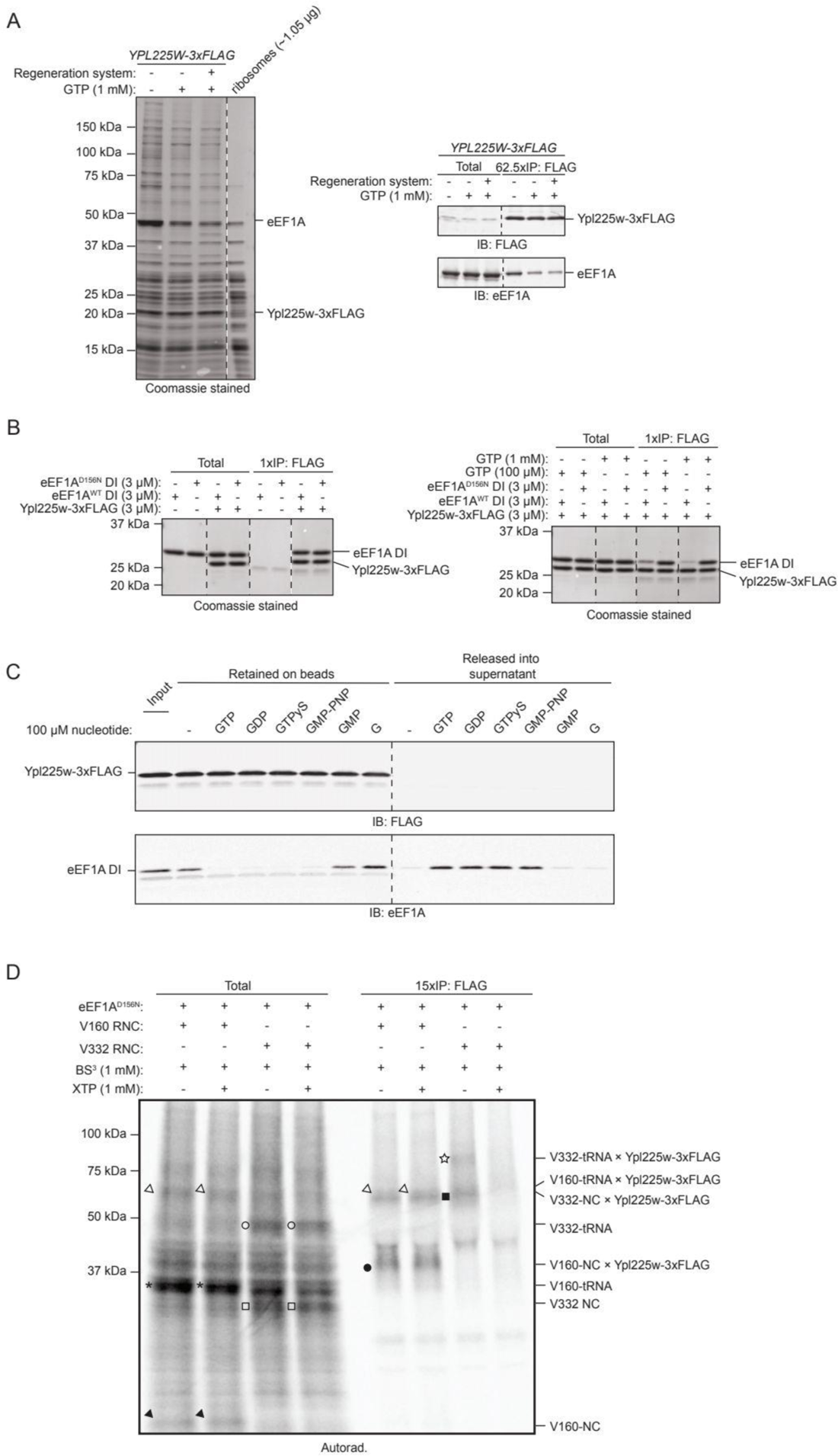
Additional evidence of GTP-dependent folding of eEF1A DI. (A) As in Fig. 5c but reactions were supplemented with GTP, or an energy regeneration system where indicated. Salt-washed ribosomes were loaded as a control. (B) As in Fig. 4c but with the indicated proteins. Dashes indicate cropping from the same Coomassie-stained gel to remove irrelevant lanes. (C) As in Fig. 6b but with 100 µM of the indicated nucleotide. (D) As in Fig. 6f but reactions were supplemented with 1 mM XTP during the full duration of translation (15 minutes) where indicated. Filled arrow: V160-NC, asterisk: V160-tRNA, filled circle: V160-NC x Ypl225w-3xFLAG, open arrow: V160-tRNA x Ypl225w-3xFLAG, open square: V332-NC, open circle: V332-tRNA, filled square: V332-NC x Ypl225w-3xFLAG, open star: V332-tRNA x Ypl225w-3xFLAG.

**Extended Data Fig. 7.**
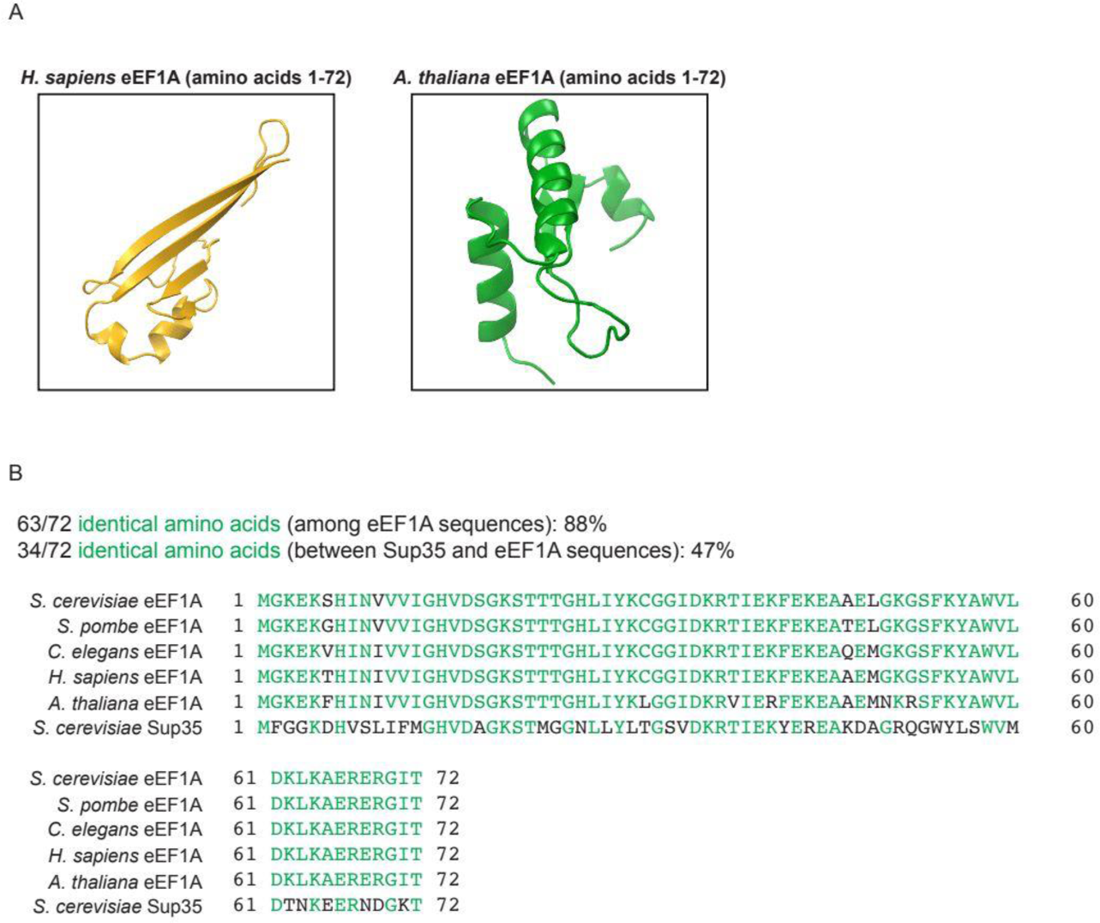
eEF1A co-translational modeling simulations reveal a correlation between eEF1A misfolding and Ypl225w presence. (A) Molecular simulations were conducted using human (*Homo sapiens*) or *Arabidopsis thaliana* eEF1A (amino acids 1-72) as described in Extended Data Fig. 1c and Methods. Snapshots of the representative minimum free-energy conformations are shown for human (yellow) and *A. thaliana* (green) eEF1A. (B) CLUSTAL Omega multiple sequence alignment of *S. pombe*, *C. elegans*, *H. sapiens*, and *A. thaliana* eEF1A along with *S. cerevisiae* eEF1A and Sup35. Identical amino acids are highlighted in green.

