## Supplementary Table 1 for "A ribosome-associating chaperone mediates GTP-driven vectorial folding of nascent eEF1A"

| Accession | WT_1 | WT_2 | WT_3 | Ypl225w_1 | Ypl225w_2 | Ypl225w_3 |
|---|---|---|---|---|---|---|
| P02994 | 6.38E+04 | 2.44E+05 | 6.01E+04 | 1.69E+07 | 2.25E+07 | 2.48E+07 |
| P11484 | 5.22E+05 | 1.82E+05 | 8.29E+05 | 8.25E+06 | 8.56E+06 | 1.23E+07 |
| P10664 | 1.56E+05 | 4.43E+05 | 3.81E+05 | 1.43E+07 | 9.16E+06 | 8.42E+06 |
| P49626 | | | 4.28E+04 | 3.38E+05 | 7.06E+05 | 2.58E+05 |
| P29453 | 7.43E+04 | 7.21E+05 | | 1.38E+07 | 1.01E+07 | 7.52E+06 |
| P14126 | 2.28E+04 | | 4.35E+04 | 5.84E+06 | 2.92E+06 | 3.72E+06 |
| P0CX38 | 9.99E+04 | 1.16E+05 | 1.40E+05 | 5.93E+06 | 7.39E+06 | 6.34E+06 |
| P0CX40 | 2.26E+04 | 7.78E+04 | 8.75E+04 | 8.83E+06 | 7.46E+06 | 7.48E+06 |
| Q08971 | 2.17E+05 | 1.29E+05 | 5.38E+04 | 3.01E+06 | 3.46E+06 | 8.08E+06 |
| P05740 | 1.61E+05 | 7.32E+04 | 1.73E+05 | 5.24E+06 | 6.44E+06 | 3.78E+06 |
| P00549 | 5.77E+05 | 1.33E+05 | 1.17E+06 | 1.83E+06 | 2.01E+06 | 1.38E+06 |
| P14127 | 5.93E+04 | | 5.62E+04 | 4.24E+06 | 5.49E+06 | 2.71E+06 |
| P33442 | 1.07E+04 | 6.00E+05 | 2.10E+05 | 4.56E+06 | 4.53E+06 | 3.69E+06 |
| P05737 | 3.39E+04 | 9.39E+04 | 9.76E+04 | 9.95E+06 | 4.99E+06 | 6.78E+06 |
| P0CX26 | 1.35E+05 | 2.89E+05 | 2.62E+05 | 3.19E+06 | 2.31E+06 | 2.45E+06 |
| P05755 | 3.95E+04 | 1.34E+05 | 9.01E+04 | 4.14E+06 | 6.80E+06 | 3.22E+06 |
| P0CX43 | | | 2.75E+04 | 1.86E+06 | 2.56E+06 | 2.50E+06 |
| P23248 | | | | 2.41E+06 | 3.20E+06 | 1.04E+06 |
| Q12213 | 9.18E+03 | | 3.07E+04 | 7.34E+05 | 1.86E+05 | 1.88E+05 |
| P39516 | 1.10E+05 | 2.63E+05 | 1.53E+05 | 5.59E+06 | 8.16E+06 | 5.82E+06 |
| P46990 | | | | 1.69E+05 | | 2.38E+05 |
| P26783 | 1.29E+05 | 1.26E+05 | 1.28E+05 | 2.75E+06 | 1.36E+06 | 5.16E+06 |
| P0CX48 | 9.73E+03 | 1.15E+06 | 7.92E+04 | 5.13E+06 | 6.63E+06 | 8.07E+06 |
| P05750 | 8.30E+03 | | | 4.24E+06 | 1.88E+06 | 2.84E+06 |
| P17076 | 2.04E+04 | | 1.97E+04 | 6.27E+05 | 2.42E+05 | 5.26E+05 |
| P0CX83 | 9.02E+04 | 3.85E+05 | 3.23E+05 | 5.55E+06 | 4.45E+06 | 6.78E+06 |
| P05748 | 2.38E+04 | 6.69E+05 | 8.60E+04 | 4.96E+06 | 7.88E+06 | 6.07E+06 |
| P06105 | | 1.08E+05 | | 1.18E+06 | 1.11E+06 | 7.45E+05 |
| P0CX46 | 4.77E+04 | | 1.12E+05 | 4.17E+06 | 5.29E+06 | 3.87E+06 |
| P53221 | 1.13E+06 | 3.28E+06 | 1.19E+06 | 9.29E+06 | 1.08E+07 | 8.48E+06 |
| Q3E754 | 4.38E+04 | | 6.68E+04 | 6.17E+04 | | 3.29E+05 |
| P0CX24 | 3.42E+04 | 1.54E+05 | 6.71E+04 | 5.02E+06 | 3.75E+06 | 5.15E+06 |
| Q12499 | 1.92E+05 | 7.87E+04 | 2.10E+05 | 6.72E+05 | 1.51E+06 | 4.70E+05 |
| P0CX85 | 5.40E+04 | 1.47E+05 | 2.80E+04 | 3.73E+06 | 1.24E+06 | 3.24E+06 |
| P05743 | 7.39E+04 | 3.44E+05 | 7.08E+04 | 5.58E+05 | 4.49E+05 | 9.78E+05 |
| P39015 | | | | 5.62E+06 | 2.11E+05 | 2.67E+06 |
| O13516 | | | | | | |
| P0C0V8 | 4.38E+04 | 7.73E+04 | 1.20E+04 | 3.48E+06 | 2.02E+06 | 3.38E+06 |
| P38701 | 4.12E+04 | 1.97E+05 | 3.56E+04 | 5.73E+06 | 4.30E+06 | 4.26E+06 |
| P00359 | 3.46E+05 | | 9.74E+05 | 2.58E+06 | 1.87E+06 | 2.11E+06 |
| P0CX31 | 3.53E+06 | 2.61E+06 | 3.13E+06 | 3.81E+06 | 4.14E+06 | 1.90E+06 |
| Q12460 | 9.02E+03 | 8.18E+04 | 3.61E+04 | 6.67E+05 | 2.45E+06 | 5.47E+05 |

| | | | | | | |
|---|---|---|---|---|---|---|
| P04449 | 2.69E+04 | | 8.04E+04 | 5.65E+06 | 6.38E+06 | 4.31E+06 |
| P26785 | | | | 4.33E+06 | 3.96E+06 | 5.56E+06 |
| P24000 | | | | | 2.05E+05 | |
| P0CX35 | | | 4.73E+04 | 4.14E+06 | 2.56E+06 | 3.45E+06 |
| P38754 | 1.94E+04 | 6.32E+05 | | 2.88E+06 | 2.12E+06 | 3.19E+06 |
| P0CX52 | | 1.05E+05 | 4.13E+04 | 2.65E+06 | 3.03E+06 | 4.68E+06 |
| P04456 | 7.94E+04 | | 8.80E+04 | 2.95E+06 | 3.07E+06 | 2.66E+06 |
| P0CX49 | 3.34E+04 | 9.36E+04 | 2.61E+04 | 4.84E+06 | 7.70E+06 | 6.03E+06 |
| P0CX53 | | | | 3.80E+06 | 7.82E+06 | 4.07E+06 |
| P38011 | 4.66E+03 | | 3.49E+04 | 2.53E+06 | 2.37E+06 | 1.38E+06 |
| P10592 | 1.52E+04 | 4.65E+05 | 6.70E+04 | 6.06E+05 | 5.40E+05 | 5.88E+05 |
| P04147 | | | 8.05E+04 | 7.03E+05 | 6.21E+05 | 6.71E+05 |
| P40212 | | 5.88E+04 | | 4.58E+06 | 7.34E+06 | 6.91E+06 |
| P19657 | 1.28E+05 | 6.17E+04 | 1.34E+05 | 7.00E+05 | 9.28E+05 | 1.15E+06 |
| P05317 | 1.64E+04 | | 5.91E+04 | 3.98E+06 | 4.03E+06 | 2.83E+06 |
| P38934 | | | 1.97E+04 | 6.25E+05 | 8.60E+05 | 1.25E+06 |
| P0C2H6 | 1.10E+04 | 8.07E+04 | 2.68E+05 | 2.30E+06 | 1.02E+06 | 1.47E+06 |
| O14455 | 2.18E+05 | 1.27E+05 | 3.62E+05 | 8.31E+06 | 7.24E+06 | 4.23E+06 |
| P10591 | | | | 3.80E+04 | | 5.20E+04 |
| P26786 | | | 7.88E+04 | 1.94E+06 | 2.92E+06 | 1.86E+06 |
| P00925 | 1.82E+05 | 9.31E+04 | 6.46E+05 | 4.46E+05 | 7.68E+05 | 6.15E+05 |
| Q3E7Y3 | 3.01E+04 | | 4.12E+04 | 8.22E+05 | 2.18E+06 | 2.02E+06 |
| P41805 | | | 2.52E+04 | 2.98E+06 | 1.21E+06 | 3.65E+06 |
| P38788 | | | | 6.11E+05 | 6.19E+05 | 4.04E+05 |
| P07280 | 5.76E+04 | 2.26E+05 | 4.71E+04 | 2.65E+06 | 1.06E+06 | 1.60E+06 |
| P32905 | 1.32E+04 | 6.15E+04 | 9.87E+04 | 1.62E+06 | 2.35E+06 | 2.47E+06 |
| P05756 | | 4.90E+04 | 1.47E+05 | 2.25E+06 | 2.37E+06 | 3.06E+06 |
| Q02326 | 1.15E+05 | | | 1.63E+06 | 1.06E+06 | 3.11E+06 |
| Q02753 | 7.55E+03 | | | 1.68E+06 | 2.15E+06 | 5.58E+06 |
| P0CX55 | 1.06E+04 | 5.97E+04 | | 1.53E+06 | 3.64E+06 | 1.43E+06 |
| P48164 | 2.95E+04 | 1.20E+05 | 2.39E+04 | 4.46E+05 | 3.37E+05 | 5.36E+05 |
| P06634 | 5.72E+04 | 8.85E+04 | | 9.25E+05 | 4.63E+05 | 6.54E+05 |
| P26784 | | 5.34E+04 | | 1.32E+06 | 4.42E+05 | 8.73E+05 |
| P05739 | 1.19E+04 | | 2.33E+04 | 1.30E+06 | 1.24E+06 | 1.29E+06 |
| P15646 | 2.73E+04 | 8.08E+04 | 6.28E+04 | 1.05E+06 | 1.03E+06 | 2.79E+05 |
| P25443 | 1.25E+04 | | 1.99E+04 | 1.74E+06 | 6.26E+05 | 1.34E+06 |
| P46784 | | | | 1.12E+06 | 1.88E+05 | 3.90E+05 |
| Q01855 | 3.25E+04 | | 8.48E+04 | 1.39E+06 | 1.03E+06 | 2.09E+06 |
| P06169 | 5.99E+04 | | | 7.44E+05 | 2.08E+05 | 8.11E+05 |
| P0C0X0 | 1.30E+05 | 1.94E+05 | 1.19E+05 | 8.57E+05 | 1.09E+06 | 6.60E+05 |
| Q3E757 | 7.83E+04 | | 5.07E+04 | 4.24E+06 | 5.41E+06 | 2.82E+06 |
| P05738 | 1.87E+04 | 2.83E+05 | 3.03E+04 | 9.89E+05 | 1.18E+06 | 1.41E+06 |
| Q99337 | | | | 3.01E+05 | 1.10E+05 | 5.08E+05 |

| | | | | | | |
|---|---|---|---|---|---|---|
| P14120 | 5.70E+04 | 8.01E+04 | 1.35E+05 | 8.89E+05 | 7.25E+05 | 8.54E+05 |
| P49167 | 3.90E+05 | 7.76E+04 | 7.28E+05 | 4.88E+05 | 2.68E+05 | 4.57E+05 |
| P32473 | | 9.36E+04 | | 4.55E+05 | 2.69E+05 | 3.84E+05 |
| P48589 | | | 2.20E+04 | 2.50E+06 | 4.69E+06 | 3.32E+06 |
| P05744 | | | | 2.11E+06 | 2.64E+06 | 6.85E+06 |
| P0CX41 | 4.49E+04 | 1.55E+05 | 4.23E+04 | 1.28E+06 | 4.35E+05 | 3.22E+05 |
| P02406 | | | 1.84E+05 | 1.25E+06 | 2.25E+06 | 1.18E+06 |
| P39939 | 2.37E+04 | | 6.71E+04 | 9.14E+05 | 9.32E+05 | 2.32E+06 |
| P02309 | | | | 3.63E+05 | 4.40E+05 | 2.34E+05 |
| P26321 | 1.01E+04 | | | 2.60E+06 | 1.19E+06 | 1.51E+06 |
| O14467 | | | | 2.47E+05 | 9.89E+05 | 5.04E+05 |
| P16521 | 4.23E+06 | 3.11E+06 | 7.08E+06 | 4.34E+05 | 6.25E+05 | 2.01E+05 |
| P32527 | | | 4.88E+04 | 1.99E+05 | 2.50E+05 | 1.37E+05 |
| P38061 | | 1.05E+05 | | 9.58E+05 | 2.26E+06 | 5.41E+05 |
| P32324 | | | | 4.53E+05 | 1.47E+04 | 4.32E+05 |
| Q3E792 | | | 2.85E+04 | 4.21E+05 | 5.30E+05 | 8.13E+05 |
| P0C2H8 | 6.01E+04 | 1.66E+05 | 1.15E+05 | 1.06E+06 | 4.17E+05 | 7.63E+05 |
| P38879 | | | | 1.30E+06 | 1.51E+06 | 4.80E+05 |
| P05319 | | | | 1.73E+06 | 1.02E+05 | 1.34E+06 |
| Q03195 | | | | 7.01E+04 | 1.03E+05 | 1.25E+05 |
| P0CX30 | | | | 2.99E+06 | 1.19E+05 | 2.23E+06 |
| P38820 | | | | 1.90E+05 | 3.15E+05 | 1.94E+05 |
| P04650 | | | 1.94E+04 | 1.97E+06 | 7.17E+05 | 2.32E+06 |
| P27476 | | 1.25E+05 | | 1.07E+05 | 1.09E+05 | 8.46E+04 |
| P40991 | | | | 1.90E+05 | 6.62E+04 | 3.11E+05 |
| P00560 | 2.79E+05 | | 4.52E+05 | 1.25E+05 | 5.84E+03 | 1.26E+05 |
| P33322 | 7.85E+05 | 1.72E+06 | 7.59E+05 | 2.27E+05 | 2.82E+05 | 3.73E+05 |
| P10622 | 1.69E+04 | | 3.45E+04 | 3.37E+05 | | 4.56E+05 |
| Q01080 | | | | 8.15E+03 | | 1.34E+05 |
| P02400 | | | | 2.41E+05 | 1.82E+05 | 1.59E+05 |
| P40433 | | | | 1.75E+05 | | 2.19E+05 |
| P40525 | 8.44E+04 | 7.02E+04 | 1.99E+05 | 2.39E+06 | 4.79E+06 | 2.08E+06 |
| P09440 | | | | 1.12E+04 | 4.45E+03 | 1.51E+05 |
| Q06697 | | | | | 2.29E+05 | |
| P16862 | 4.05E+04 | | | | 1.09E+05 | |
| P16861 | | | | 1.12E+05 | | 1.90E+05 |
| P89105 | | | | 8.79E+04 | | 1.64E+05 |
| P12695 | 2.99E+04 | 7.60E+04 | | 3.15E+04 | 5.14E+04 | 1.28E+05 |
| P51402 | | | | 5.98E+05 | 1.02E+05 | 9.57E+05 |
| Q12754 | | | | | 1.57E+05 | |
| P38439 | | | | 3.12E+04 | 2.14E+05 | 2.45E+04 |
| P12385 | | | | 5.50E+04 | 9.01E+03 | 6.54E+04 |
| P53064 | | | | 2.91E+04 | 5.97E+04 | 3.45E+04 |

| | | | | | | |
|---|---|---|---|---|---|---|
| P47006 | | | | 2.25E+04 | 6.20E+04 | 6.34E+04 |
| Q12159 | 1.46E+04 | | 3.38E+04 | | 2.80E+05 | |
| P53297 | | | | 3.02E+04 | 2.73E+04 | 4.71E+04 |
| P05759 | | | | 5.05E+05 | 1.72E+06 | 2.06E+05 |
| P36049 | | | | | 7.63E+04 | |
| P49166 | | | | 4.37E+05 | 1.75E+05 | 1.72E+05 |
| P09624 | | | | 2.97E+03 | 4.71E+04 | |
| Q08237 | | | | 9.15E+03 | 8.93E+04 | 1.92E+04 |
| P0CX27 | | | | 1.19E+06 | 3.72E+05 | 4.22E+05 |
| P28000 | | | | | | |
| P40961 | | | | | 3.13E+04 | |
| Q08208 | | | | | 6.49E+04 | |
| P25567 | 1.25E+04 | 5.40E+04 | 3.87E+04 | | 2.31E+04 | 5.15E+03 |
| P41058 | | 7.47E+04 | | 4.46E+05 | 1.07E+05 | 2.89E+05 |
| P41057 | | 1.03E+05 | | 6.13E+04 | 2.23E+05 | 1.49E+05 |
| P33421 | | | | | 3.28E+05 | |
| P07260 | | | | 2.04E+04 | | 5.62E+04 |
| P02381 | | | | | | |
| P38911 | 1.44E+04 | 8.04E+04 | | 1.33E+05 | 1.98E+05 | 5.97E+04 |
| P04912 | | | | | 4.41E+05 | |
| Q05022 | | | | 3.84E+04 | 2.31E+05 | 6.12E+04 |
| P0CH09 | | | | 1.08E+05 | | 1.00E+05 |
| P38711 | | | | 1.42E+06 | 6.39E+05 | 4.41E+05 |
| P39990 | | | | 2.76E+05 | 5.87E+05 | 5.86E+04 |
| P00931 | | | 5.62E+04 | 5.10E+05 | 1.28E+06 | 4.39E+05 |
| Q07362 | | | | 3.48E+04 | | 7.87E+04 |
| Q03532 | | | | | | |
| P22138 | | | | 4.68E+04 | | 2.78E+04 |
| P33201 | | | | 1.14E+04 | 2.67E+05 | |
| P53914 | | | | | | |
| Q08287 | | | | | | |
| Q06506 | | | | | | |
| P50095 | | | | 7.01E+03 | | 1.17E+05 |
| P0CS90 | | | | | | |
| P19882 | | | | 9.16E+03 | | 1.06E+04 |
| Q06108 | | | | 2.53E+05 | | 8.06E+04 |
| P40988 | | | | | | |
| P38110 | | 5.92E+04 | | 3.91E+05 | 6.48E+05 | 4.89E+05 |
| P39730 | | | | | 5.82E+04 | |
| P10080 | | | | | | |
| P40010 | | | | | 1.39E+04 | |
| P39935 | | | | 1.67E+04 | | 3.26E+04 |
| P60010 | | | | 8.97E+04 | | 2.11E+04 |

| | | | | | | |
|---|---|---|---|---|---|---|
| P02293 | | | | | 2.96E+05 | |
| Q04471 | | | | 2.14E+04 | 6.79E+04 | 1.07E+04 |
| P17255 | | 1.30E+05 | | 5.24E+03 | 1.29E+04 | 4.70E+04 |
| P40466 | 5.24E+06 | 7.49E+04 | | | | 1.37E+04 |
| P40850 | | | | 2.26E+04 | | 1.28E+04 |
| Q12143 | | | | | | |
| P61830 | | | | 6.51E+04 | | 7.34E+04 |
| P12753 | 5.71E+06 | 4.86E+06 | 5.80E+06 | | 1.34E+05 | 1.25E+04 |
| P00950 | | | | 3.69E+04 | 2.77E+05 | 2.10E+04 |
| P38261 | | | | | 9.27E+04 | |
| Q12000 | | | | | 1.10E+05 | |
| Q12180 | | 4.84E+05 | | | 1.01E+05 | |
| P05747 | | 5.04E+04 | | | 2.09E+05 | 1.97E+05 |
| P40693 | | | | 1.82E+04 | 3.79E+04 | |
| P32583 | | | | 1.86E+04 | | 1.34E+04 |
| Q06339 | | | | | | |
| P08964 | | | | 3.00E+04 | | 1.08E+05 |
| P00927 | | | | 1.87E+06 | | 4.35E+06 |
| Q12136 | | | | | | |
| Q03690 | | | | | | |
| Q02948 | | | | 6.89E+02 | 9.75E+04 | |
| P16387 | | | | | 6.17E+04 | |

WT = Untagged (*YPL225W)*

Ypl225w = (*YPL225W-3xFLAG)*

The numbers next to the indicated strain represent the replicate.
