## Supplementary Tables 2-4 for "A ribosome-associating chaperone mediates GTP-driven vectorial folding of nascent eEF1A"

**Supplementary Table 1.** **Ypl225w-3xFLAG IP-MS results.**

### Supplementary Table 2: Yeast strains

| Yeast Strains | | |
|---|---|---|
| *W303 MATa leu2-3,112::4xHSE-YFP::CgLEU2 trp1-1 can1-100 ura3-1 ade2-1 his3-11,15* | Zheng et al. (2016) | VDY3334 |
| *W303 MATa leu2-3,112::4xHSE-YFP::CgLEU2 trp1-1 can1-100 ura3-1 ade2-1 his3-11,15 ypl225wΔ::hphMX6* | This study | VDY6165 |
| *W303 MATa leu2-3,112::4xHSE-YFP::CgLEU2 trp1-1 can1-100 ura3-1 ade2-1 his3-11,15 YPL225W-3xFLAG::kanMX* | This study | VDY6217 |
| *W303 MATa leu2-3,112::4xHSE-YFP::CgLEU2 trp1-1 can1-100 ura3-1 ade2-1 his3-11,15 YPL225W-3xFLAG::kanMX EGD2-13xMyc::HIS3* | This study | VDY6292 |
| *W303 MATa leu2-3,112 trp1-1 can1-100 ura3-1 ade2-1 his3-11,15* | Solís et al. (2016) | W303 (VDY465) |
| *W303 MATa leu2-3,112 trp1-1 can1-100 ura3-1 ade2-1 his3-11,15 ypl225wΔ::HIS3 ura3-1::prGAL1-eEF1A-DI-GFP::URA3* | This study | VDY6205 |
| *W303 MATa leu2-3,112::pZ4EV-OsTIR1(F74G)_pACT1-Z4Ev-ATF_CgLEU2 trp1-1 can1-100 ura3-1 ade2-1 his3-11,15 TEF1-yeGFP::hphMX* | Sabbarini et al. (2023) | VDY6176 |
| *W303 MATa leu2-3,112::pZ4EV-OsTIR1(F74G)_pACT1-Z4Ev-ATF_CgLEU2 trp1-1 can1-100 ura3-1 ade2-1 his3-11,15 TEF1-yeGFP::hphMX ypl225wΔ::HIS3* | This study | VDY6206 |
| *W303 MATa leu2-3,112::4xHSE-YFP::CgLEU2 trp1-1 can1-100 ura3-1 ade2-1 his3-11,15 ypl225wΔ::hphMX6 EGD2-13xMyc::HIS3* | This study | VDY6334 |
| *W303 MATa leu2-3,112::4xHSE-YFP::CgLEU2 trp1-1 can1-100 ura3-1 ade2-1 his3-11,15 YPL225W*$^{F19A}$*-3xFLAG::kanMX EGD2-13xMyc::HIS3* | This study | VDY6341 |
| *W303 MATa leu2-3,112::4xHSE-YFP::CgLEU2 trp1-1 can1-100 ura3-1 ade2-1 his3-11,15 hsf1Δ::kanMX pNH604-PrHSF1-HSF1(1–424)::TRP1* | This study | VDY6335 |
| *W303 MATa leu2-3,112::4xHSE-YFP::CgLEU2 trp1-1 can1-100 ura3-1 ade2-1 his3-11,15 hsf1Δ::kanMX pNH604-PrHSF1-HSF1(1–424)::TRP1 ypl225wΔ::hphMX6* | This study | VDY6285 |
| *W303 MATa leu2-3,112::4xHSE-YFP::CgLEU2 trp1-1 can1-100 ura3-1 ade2-1 his3-11,15 hsf1Δ::kanMX pNH604-PrHSF1-HSF1::TRP1* | This study | VDY6336 |
| *W303 MATa leu2-3,112::4xHSE-YFP::CgLEU2 trp1-1 can1-100 ura3-1 ade2-1 his3-11,15 hsf1Δ::kanMX pNH604-PrHSF1-HSF1::TRP1 ypl225wΔ* | This study | VDY6287 |
| *W303 MATa leu2-3,112::4xHSE-YFP::CgLEU2 trp1-1 can1-100 ura3-1 ade2-1 his3-11,15 YPL225W-3xFLAG::kanMX egd2Δ::hphMX6* | This study | VDY6283 |
| *W303 MATa ura3-1::4xHSE-GFP:URA3 TRP1::prGAL1-TEF2 HIS3::prGAL1-TEF1 can1-100 ade2-1 YPL225W-3xFLAG::kanMX* | This study | VDY6238 |
| *W303 MATa leu2-3,112 trp1-1 can1-100 ura3-1 ade2-1 his3-11,15 ypl225wΔ::hphMX6* | This study | VDY6374 |
| *W303 MATa leu2-3,112 trp1-1 can1-100 ura3-1 ade2-1 his3-11,15* | This study | VDY6375 |

### Supplementary Table 3: Plasmids

| Plasmids | | |
|---|---|---|
| pET29b-eEF1A-DI-myc-6xHis | This study | pVD2918 |
| pET29b-eEF1A-DI (D156N)-myc-6xHis | This study | pVD2935 |
| pET29b-Sup35-DI-myc-6xHis | This study | pVD2995 |
| pET29b-Ypl225w-3xFLAG-3c-6xHis | This study | pVD2936 |
| pET29b-Ypl225w (F19A)-3xFLAG-3c-6xHis | This study | pVD2938 |
| pETDuet-myc-EGD2_10xHis-EGD1 | This study | pVD2926 |
| pETDuet-myc-EGD2 (ΔUBA)_10xHis-EGD1 | This study | pVD2927 |
| pETDuet-myc-EGD2 (L165A)_10xHis-EGD1 | This study | pVD2967 |
| pETDuet-myc-EGD2 (V166A)_10xHis-EGD1 | This study | pVD2968 |
| pETDuet-myc-EGD2 (L165AV166A)_10xHis-EGD1 | This study | pVD2969 |
| pET16b-10xHis-3c-3xFLAG-eEF1A | Sabbarini et al. (2023) | pVD2609 |
| *pmGCN4-EGFP* | Gift from Onn Brandman | pVD2746 |
| pRS414: *prTEF2-TEF2*$^{D156N}$ | McQuown et al. (2023) | pVD2860 |
| pRS406:*prGAL1-eEF1A DI-GFP* | Sabbarini et al. (2023) | pVD2659 |
| *pRS414:TEF2pr-eEF1A DI-GFP* | This study | pVD3006 |
| *pRS414:TEF2pr-SUP35 DI-GFP* | This study | pVD3007 |

### Supplementary Table 4: Oligos

| Oligonucleotides | | |
|---|---|---|
| GGGGGTAATACGACTCACTATAGGGAGAatttcatacacaata<br>taaacgattgccaccATGGGTA<br>AAGAGAAGTCTCAC | Sabbarini et al. (2023) | oVD13102 |
| TTTTTTTTTTTTTTTTTTTTttacat ctacactgttgttatcagtcgggcTCATTT<br>CTTAGCAGCCTTTTGAGCAGCC | Sabbarini et al. (2023) | oVD13103 |
| AACTTTGACGGAGTCCATCTTGTTGAC | This study | oVD13432 |
| AACTGGATCGTTCTTAGCGTCACC | This study | oVD13603 |
